# Microbial valerate is associated with CAR T dysbiosis and its supplementation enhances CAR T function in B-cell lymphoma

**DOI:** 10.64898/2026.09.01.748711

**Authors:** Lubna Rehman, Wenting Song, Abdur Rehman, Kamini Singh, Shiv Govind Rawat, Faezeh Darbaniyan, Cao Cuong Le, Federico Mario Aletti, Jinsheng Weng, Jingwei Liu, Xiayoun Cheng, Yongfu Tang, Sridevi Patchva, Matthew D. Richard, Fuliang Chu, Jingjing Cao, Christopher R. Flowers, Elizabeth J. Shpall, Mark R. Tanner, Johannes Fahrmann, Eran Elinav, Christoph K. Stein-Thoeringer, Michael D. Jain, Robert R. Jenq, Abhinav Jain, Sattva S. Neelapu, Ye Zheng, Neeraj Y Saini

## Abstract

**Key Points:**

- Anaerobe-targeting antibiotic exposure marks low gut valerate and apheresis CAR-T products with effector dysfunction.
- Valerate supplementation rescues CAR-T deficits in dysbiotic hosts, identifying valerate as an actionable biomarker and target.

Anaerobe-depleting antibiotic exposure is associated with inferior progression-free survival after CD19 CAR T-cell therapy in large B-cell lymphoma, yet the cellular mechanisms linking gut dysbiosis to the CAR T-cell product and whether this imprint is reversible have remained undefined. In two independent CAR-T candidate cohorts, low stool valerate at the time of CAR-T eligibility identified a multi-metabolite-deficient dysbiotic gut microbiome state marked by depletion of fiber-fermenting commensals and loss of carbohydrate-fermentation, SCFA-biosynthesis, and amino-acid metabolism pathways. Reanalysis of single-cell RNA sequencing from 42 lymphoma patients stratified by piperacillin-tazobactam/imipenem/meropenem (PIM) exposure revealed that PIM-exposed CAR T-cell products were CD4-skewed, with significantly elevated AP-1/immediate-early gene (IEG) and cellular activation signatures that together predicted inferior progression-free survival. Ex vivo conditioning of CAR T-cells with valerate produced a chromatin and transcription factor program distinct from butyrate or propionate, characterized by KLF/SP/EGR family engagement, KLF4 promoter opening, and broad induction of AP-1/IEG and MHC class II transcripts, whereas butyrate drove broader chromatin remodeling with TBX21/EOMES/NF-κB gains and KLF2 promoter closure, and propionate induced an NFY-centered program with preferential commitment to low-mitochondrial-content states. Untargeted metabolomics confirmed valerate uptake and mitochondrial β-oxidation in CAR T-cells, while dietary sodium valerate supplementation in meropenem-treated mice bearing A20 lymphoma significantly reduced tumor burden and extended survival compared with CAR T-cells alone. These findings identify stool valerate as a bedside-deployable biomarker of dysbiosis-imprinted CAR T-cell dysfunction and support ex vivo or dietary valerate supplementation as a clinically tractable strategy to improve CAR-T anti-tumor function in patients with disrupted gut microbiomes.

## Introduction

The gut microbiome has emerged as an important upstream modifier of clinical outcomes across immune-checkpoint therapy^1^, allogeneic hematopoietic stem-cell transplantation^2^, and most recently chimeric antigen receptor (CAR) T-cell therapy^3^. In patients with large B-cell lymphoma (LBCL), microbiome dysbiosis often develops from multiple converging upstream factors, including antibiotic exposure, cancer cachexia with reduced dietary fiber intake, prior chemotherapy damage to gut mucosa, and concomitant proton-pump inhibitors or steroids, among others^4–8^. These insults collectively deplete obligate-anaerobic gut commensals and reduce the microbial metabolite output that supports systemic immune function.

We and others have previously reported across two independent international cohorts that pre-CAR T-cell exposure to broad-spectrum antibiotics targeting obligate anaerobes is associated with inferior progression-free survival (PFS) in LBCL patients^3,9,10^. This exposure identifies a high-risk patient population with markedly depleted levels of short-chain fatty acids (SCFAs), particularly valeric acid (VA, –pentanoic acid), and other immunomodulatory microbial metabolites in both stool and plasma^10^. However, the cellular and molecular link between this dysbiotic gut state and the CAR T-cell product itself has not been defined.

SCFAs produced by anaerobic fiber fermentation are the most extensively characterized class of immunomodulatory microbial metabolites and enhance CD8 T-cell antitumor function in preclinical adoptive cell therapy models^11–14^. Among the four major SCFAs, VA is distinguished by its potent dual class I/II HDAC inhibition, acetyl-CoA-pool feeding through β-oxidation, and entry into the TCA cycle through both acetyl-CoA and succinyl-CoA^15,16^. These characteristics establish valerate as a metabolite of particular interest for cellular therapy. Although these properties have been described in healthy donor T-cells and murine models, it remains unknown whether valerate directly alters CAR T-cells, including how it affects these cells at the chromatin and transcription factor level. Furthermore, it is unknown whether valerate supplementation can rescue microbiome-metabolite defects imprinted on patient-derived CAR T-cell apheresis products, or whether restoring valerate in a dysbiotic host improves antitumor activity in vivo. Critically, no clinically actionable biomarker has been linked to these cellular phenotypes.

Here, across two independent CAR T-cell candidate cohorts at the University of Texas MD Anderson Cancer Center (MDACC) and Germany, we show that low stool valerate marks a reproducible, multi-metabolite-deficient gut state characterized by loss of fiber-fermenting commensals and coordinated reduction in SCFAs, B-vitamin, and amino acid biosynthetic pathways. This dysbiotic state imprints a distinct, prognostically adverse cellular phenotype on the apheresis CAR T-cell product. Through integrated single-cell RNA-seq (scRNA-seq), ATAC-seq, regulon, metabolomic, and in vivo rescue experiments, we establish that valerate supplementation, either during ex vivo manufacturing or as a dietary intervention, partially corrects these deficits and improves CAR T-cell antitumor function. Together, these findings indicate stool valerate as a candidate companion diagnostic and identify two complementary intervention strategies, manufacturing-stage conditioning and dietary host-side supplementation, for improving CAR T-cell function in microbiome-disrupted LBCL patients.

## Methods

### Clinical cohort

The clinical cohort consisted of adult patients with relapsed or refractory LBCL treated with anti-CD19 CAR T-cell therapy at MDACC (USA discovery cohort)^10^. A German validation cohort of patients treated at the hospital of the Ludwig Maximilians Universität Munich and Heidelberg University Hospital was used for independent cross-cohort replication. The two cohorts were similar in clinical composition to those described in our previous reports^3,10^. All patients provided written informed consent, and the study was conducted under relevant institutional review board-approved protocols.

All other methods, including gut microbiome profiling, HUMAnN3 functional pathway analysis, single-cell RNA sequencing of apheresis products, ex vivo CAR T-cell manufacturing and conditioning with SCFAs, ATAC-seq, transcription factor regulon analysis, intracellular metabolomics, and in vivo mouse experiments, are provided in the Supplement.

## Results

### Low stool valerate identifies a multi-metabolite-deficient gut state across two independent cohorts

We previously found that stool VA levels at the time of therapy infusion independently predict PFS in LBCL patients treated with anti-CD19 CAR T-cell therapy^10^. To understand the microbial and metabolic features behind this biomarker, we reanalyzed our previous data and characterized the gut microbiome and functional pathways through whole shotgun genomic sequencing in two independent cohorts: a USA discovery cohort (n=69) and a German validation cohort (n=54). Detailed patient demographic and clinical characteristics are provided in **Supplementary Table S1**.

Stool VA levels were positively correlated with fiber-fermenting bacteria from the Lachnospiraceae and Oscillospiraceae families and negatively correlated with *Enterococcus faecium* in both cohorts (**Figure 1A**). *Anaerostipes hadrus*, *Faecalibacterium prausnitzii*, *Bifidobacterium longum*, *Fusicatenibacter saccharivorans*, *Blautia obeum*, *Roseburia hominis*, *Roseburia intestinalis*, and *Coprococcus comes* were the leading positive species in the discovery cohort (p<0.001 for the top hits). Eleven of the 13 top valerate-associated species in the discovery cohort showed the same direction in the validation cohort, with eight independently reaching statistical significance in both cohorts (**Figure 1B**). *Enterococcus faecium* replicated as the dominant inverse marker in the validation cohort (p<10⁻⁶).

**Figure 1.**
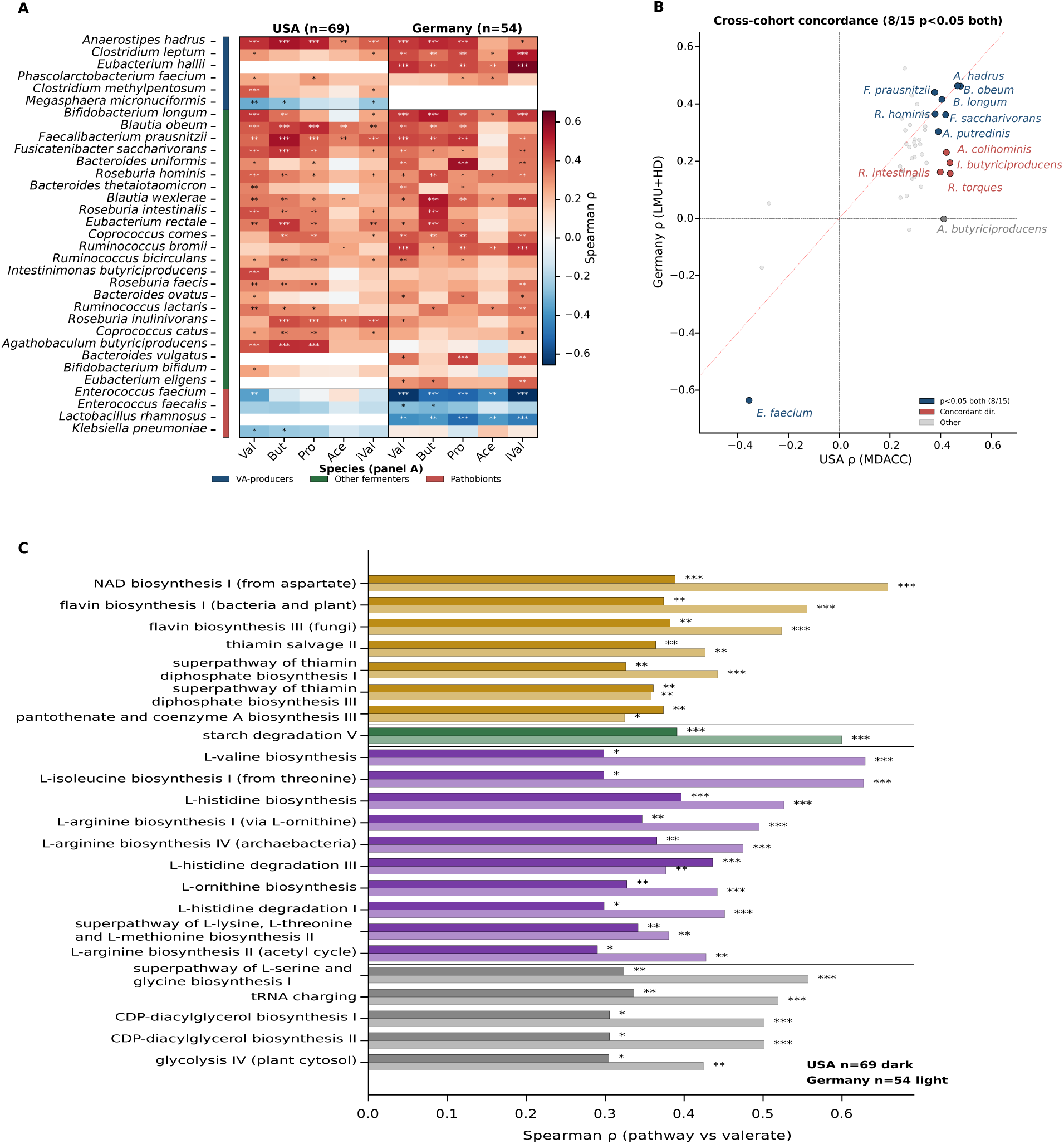
Stool valerate at CAR T-cell infusion indicates a fiber-fermenting commensal community and its multi-metabolite functional capacity, replicating in independent USA (discovery) and German (validation) cohorts. Profiling of stool SCFA and shotgun metagenomics was performed in 69 patients from MD Anderson Cancer Center (USA discovery cohort, MetaPhlAn4) and 54 patients from LMU Munich and Heidelberg (German validation cohort, MetaPhlAn3). **(A)** Spearman correlation of stool SCFAs with bacterial species in USA (left) and German (right) cohorts. The 33 species reaching p < 0.05 against stool valerate in either cohort are shown, grouped into three categories by left side-bar (canonical VA-producers, blue; other obligate-anaerobic fermenters, green; pathobionts, red). Val, valerate; But, butyrate; Pro, propionate; Ace, acetate; iVal, isovalerate. **(B)** Cross-cohort concordance of the top 15 MDACC valerate-associated species, USA ρ versus German ρ. Blue, p < 0.05 in both cohorts; red, concordant direction only; gray, other species. Red dotted line, identity. **(C)** HUMAnN3 pathways reaching BH-FDR < 0.10 in both cohorts independently with concordant direction (n = 23), paired bars for USA (solid) and Germany (faded), grouped by functional family. * p < 0.05*, ** p < 0.01, *** p < 0.001*.

We also performed functional pathway analysis using HUMAnN3^17^. Eighty-four of 528 detected pathways correlated with stool valerate at a false-discovery rate (FDR) < 0.10 in the discovery cohort, and 23 of these replicated with the same direction in the validation cohort (**Figure 1C**). The most consistent pathways involved bacterial cofactor and B-vitamin biosynthesis, including de novo NAD synthesis from aspartate (p < 0.001 in both cohorts), flavin biosynthesis, thiamin salvage, and pantothenate biosynthesis. Branched-chain amino acid pathways (valine and isoleucine) and basic amino acid pathways (arginine, ornithine, histidine) also replicated, as did starch degradation pathways. Butyrate fermentation pathways were significant in the discovery cohort and trended the same way in the validation cohort but did not pass FDR correction. Seventeen of 23 cross-cohort replicating pathways increased steadily across valerate tertiles (**Supplementary Figure S1**).

Reduced stool VA is therefore an indicator of a multi-metabolite-deficient gut microbiome state with coordinated reductions across SCFA biosynthesis, fiber utilization, branched-chain amino acid biosynthesis, and cofactor pathways. We next examined whether low valerate imprints a distinct cellular phenotype on the apheresis CAR T-cell product.

### Anaerobe-depleted gut metabolic state imprints a CD4-skewed chronic-activation CAR T-cell phenotype that predicts inferior outcomes

To determine whether this dysbiotic gut state influences the CAR T-cell product, we reanalyzed scRNA-seq data, published by Deng et al, of CAR T-cell infusion product samples of 42 LBCL patients at MDACC who received axicabtagene ciloleucel^18^. We previously reported that stool valerate is reduced at the time of CAR T-cell infusion after anaerobe-active antibiotic exposure, with the largest reductions seen in patients exposed to piperacillin-tazobactam, imipenem, or meropenem (PIM)^10^. We stratified patients by PIM exposure (8 exposed and 34 unexposed) and analyzed 164,114 CD4 and 185,226 CD8 T-cells by scRNA-seq.

Cell subsets that predicted favorable CAR T-cell outcomes in Deng et al.^18^ were depleted in the apheresis products of PIM-exposed patients (**Figure 2A and Supplementary Table S2**). The activated effector CD8 cluster was 50-fold lower (log2FC=−5.65, p=0.063), and the effector memory CD8 cluster was 11-fold lower (log2FC=−3.42) in the apheresis products of PIM-exposed versus PIM-unexposed patients. At the compartment level, the CD8 T-cell fraction fell from 48% to 32% (p=0.016) and the CD4 fraction rose from 38% to 56% (p=0.022, **Figure 2B**) in apheresis products from the PIM-exposed compared to PIM-unexposed groups. This CD4-skewed phenotype is a known predictor of inferior CD19 CAR T-cell outcomes^18,19^. Several T-cell subsets also trended higher in PIM-exposed apheresis products, including TH17 (log2FC=+0.59), Treg (log2FC=+1.17), and IFN-activated CD8 (IACs, log2FC=+2.33).

**Figure 2.**
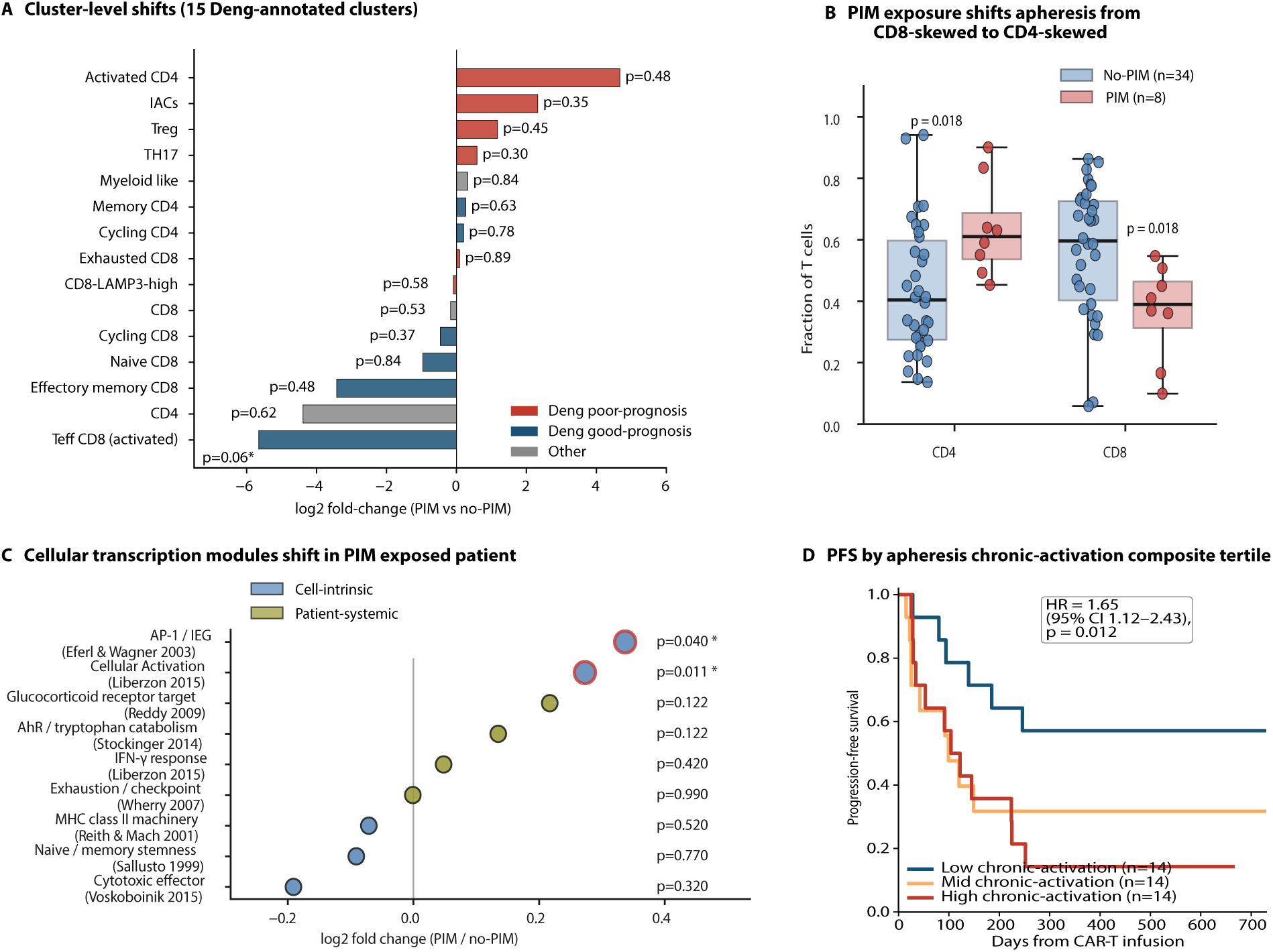
Anaerobe-active antibiotic exposure imprints the apheresis CAR T-cell product with a CD4-skewed, AP-1-activated phenotype that predicts inferior PFS. Apheresis CAR-T scRNA-seq from 42 lymphoma patients (Deng et al., Nat Med 2020) stratified by 30-day pre-apheresis piperacillin-tazobactam, imipenem, or meropenem (PIM) exposure (n = 8 exposed, 34 unexposed). **(A)** Log2 fold-change in cluster proportion in PIM-exposed vs unexposed CAR T-cell apheresis products across 15 Deng-annotated clusters (n = 5 PIM, 19 no-PIM with cluster-resolution annotation), colored by Deng’s published prognostic class (blue, good prognosis; red, poor prognosis; grey, other). Per-cluster patient-level Mann-Whitney p-values shown. Nine of 12 prognostically classified clusters shift in the hypothesized direction (binomial p = 0.073). **(B)** CD4 and CD8 compartment fractions in PIM-exposed (red, n = 8) vs unexposed (blue, n = 34) CAR T-cell apheresis products. Boxplots show median and IQR; individual patients overlaid. Patient-level Mann-Whitney. **(C)** Per-patient median scores for nine published T-cell-state signatures (Scanpy score_genes; compositions in Supplementary Methods), tested in PIM-exposed vs unexposed apheresis by patient-level Mann-Whitney. Each signature is labeled with its original publication. Points colored by category (blue, cell-intrinsic; olive, patient-systemic). Red outline = raw p < 0.05. **(D)** Kaplan-Meier PFS stratified by tertile of the apheresis chronic-activation composite score (z-summed AP-1/IEG + hallmark TNF-α/NF-κB signatures). Insert shows Cox hazard ratio per tertile with 95% CI and p-value.

To characterize the transcriptional state of the CAR T-cell infusion product, we tested a library of pre-published^20^, literature-defined T-cell state signatures between PIM-exposed and unexposed apheresis products. The AP-1/immediate-early-gene (IEG) signature (FOS, JUN, FOSB, EGR1^21,22^, log2FC=+0.34, p=0.040) and the broader cellular activation signature (MSigDB hallmark^23^, mean 0.62 vs 0.51; p=0.011) were elevated in the PIM-exposed apheresis products (**Figure 2C**). In contrast, the cytotoxic effector signature (GZMA, GZMB, PRF1, NKG7, GNLY^24^), MHC class II machinery signature (HLA-D family + CIITA + RFX trimer^25^), and naive/memory signature (TCF7, LEF1, IL7R, CCR7, SELL^26,27^) did not differ significantly between groups. Together, these shifts suggest an activation pattern in which the AP-1/IEG transcripts are elevated without parallel induction of MHC class II and cytotoxic effector programs. Patient-systemic signatures including the glucocorticoid receptor target signature (FKBP5, NR3C1, TSC22D3, SGK1, KLF9^28^), aryl hydrocarbon receptor/tryptophan catabolism signature (CYP1A1, CYP1B1, IDO1^29^), and the hallmark IFN-γ response^23^ trended upward in PIM-exposed apheresis products, albeit without statistical significance. Together, these findings describe a chronically stressed phenotype in the CAR T-cell infusion product. We infer this is imprinted by the dysbiotic gut, depleted SCFAs, and chronic systemic inflammation in PIM-exposed patients^3,9,10^, reflecting pre-manufacturing T-cell dysfunction that persists through CAR T-cell manufacturing.

The PIM-imprinted CAR-T cellular state was prognostic of PFS. In Cox regression analysis across all 42 patients, the AP-1/IEG module predicted PFS (HR=1.75, 95% CI 1.23-2.49, p=0.002), as did JUN (HR = 1.80), FOS (HR = 1.63), PDCD1 (HR=1.57, p=0.008), and the exhaustion module (HR=1.50, p=0.046). On multivariable analysis, the composite chronic-activation score remained associated with PFS (HR=1.62, 95% CI 1.06-2.48, p=0.027, **Figure 2D and Supplementary Figure S3**). Thus, the low-valerate gut microbiome state leaves a measurable, prognostically relevant imprint on the CAR T-cell product.

### Valerate, butyrate, and propionate engage three distinct CAR T-cell-state programs

We next asked whether ex vivo SCFA conditioning during CAR T-cell manufacturing could shift the phenotype of the final CAR T-cell product, and whether three particularly abundant colonic SCFAs differentially alter this phenotype. Anti-CD19-CAR T-cells from a healthy donor were manufactured using the axicabtagene ciloleucel (AXI) construct and conditioned during ex vivo expansion with culture media alone (AXI), or 1 mM butyric acid (BA), propionic acid (PA), or VA (**Figure 3A**). Expanded products were profiled by scRNA-seq (38,476 passing quality control; three replicates per condition). Major lineages were annotated using canonical markers. Within each lineage, transcriptionally distinct subclusters were defined by a combined supervised and unsupervised annotation strategy, yielding 18 CD4⁺ and 14 CD8⁺ subclusters (**Figure 3B, C**).

**Figure 3.**
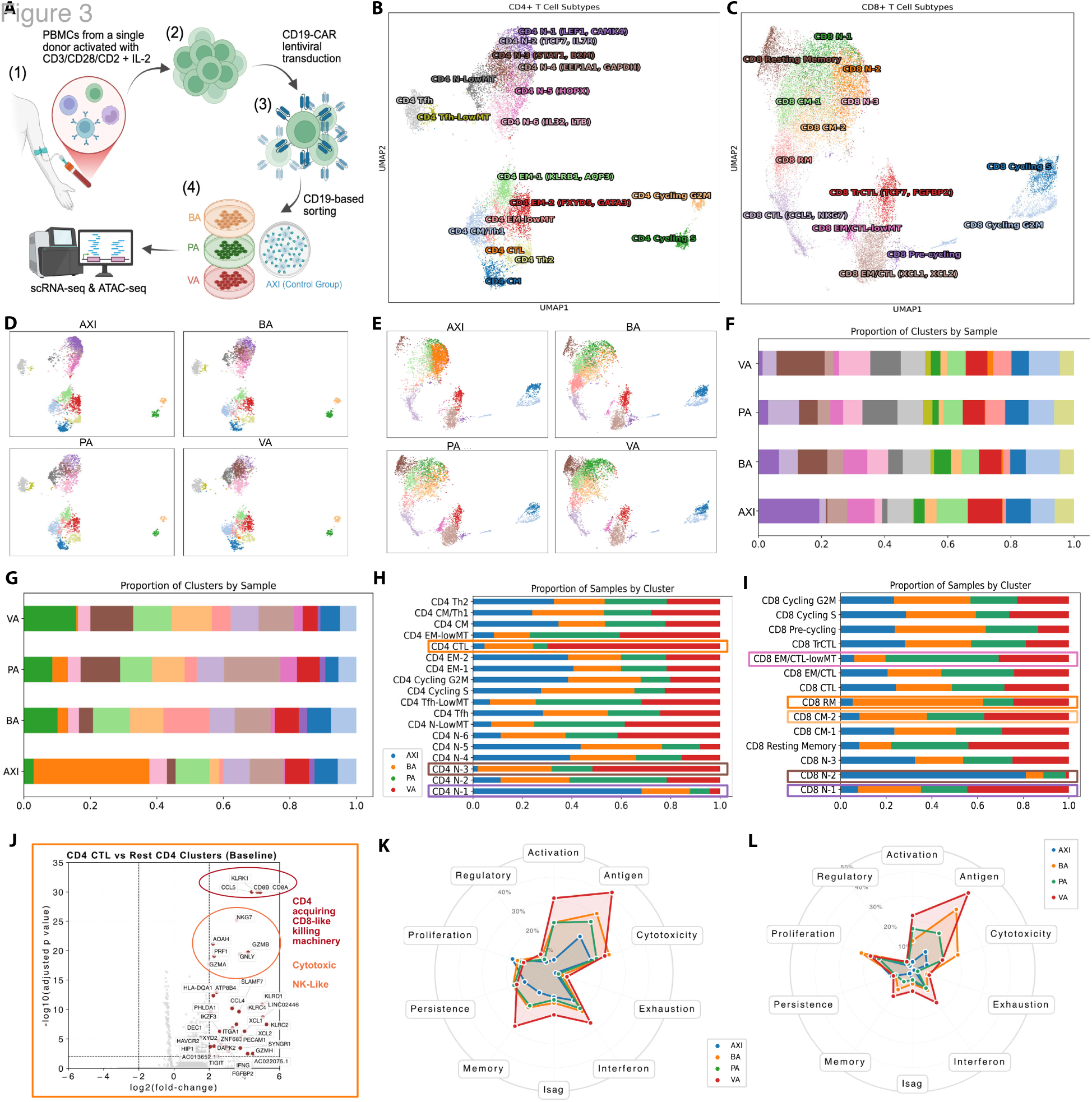
Valerate, butyrate, and propionate engage three mechanistically distinct CAR T-cell-state reprogramming programs at equimolar 1 mM ex vivo conditioning. CD19-CAR T-cells from a healthy donor were conditioned during ex vivo expansion with vehicle (AXI), butyric acid (BA), propionic acid (PA), or valeric acid (VA) at 1 mM and profiled by single-cell RNA-seq (39,739 cells, three replicates per condition). **(A)** Experimental schema. **(B, C)** UMAP embeddings of the CD4⁺ **(B)** and CD8⁺ **(C)** CAR T-cell compartments annotated into transcriptionally distinct subclusters. **(D, E)** Per-condition UMAPs of CD4⁺ **(D)** and CD8⁺ **(E)** cells showing SCFA-graded compositional shifts with AXI and VA at opposite ends and BA and PA intermediate. **(F, G)** Proportion of CD4⁺ **(F)** and CD8⁺ **(G)** subclusters per condition. **(H, I)** Proportion of CD4⁺ **(H)** and CD8⁺ **(I)** samples per cluster, with boxed clusters highlighting AXI-enriched quiescent naïve states (CD4 N-1/N-2, CD8 N-2), VA-enriched activated-naïve and cytotoxic/resident memory states (CD4 N-3, CD4 CTL, CD8 N-1, CD8 RM, CD8 CM-2), and BA-enriched effector memory cytotoxic state (CD8 EM/CTL-lowMT). **(J)** Volcano plot of CD4 CTL vs rest of CD4 cells showing canonical cytotoxic effectors plus CD8-lineage cytotoxic machinery. **(K, L)** Radar plots of CD4⁺ **(K)** and CD8⁺ **(L)** functional axis scoring across nine canonical T-cell programs. Color coding: AXI blue, BA orange, PA green, VA red. Granular per-cluster characterization is provided in the Supplement.

Both the CD4 and CD8 compartments shifted in an SCFA-graded manner with AXI and VA at opposite ends of the embedding, while BA and PA exhibited intermediate but distinct positions (**Figure 3D, E**). These shifts produced reproducible changes in the per-condition subcluster composition (**Figure 3F, G**). At the cluster-by-cluster level (**Figure 3H-L**), AXI-treated cells preferentially populated a quiescent TCF7-high naïve cluster in each compartment (CD4 N-1, 19% with AXI vs 1.3% under VA, log2FC=−3.9; CD8 N-2, 35% of CD8 cells with AXI vs 0.5% under VA, log2FC=−6.0). All three SCFAs shifted cells out of these dominant naïve populations, but the destination cell states differed by SCFA.

VA redistributed CD8 cells from the naïve compartment into multiple memory-associated clusters: CD8 N-1 (5.3-fold expansion vs AXI), CD8 resting memory (4.9-fold), CD8 effector memory (EM)/CTL-low mitochondrial content (lowMT) (4.6-fold), and CD8 CM-2 (4.1-fold). In the CD4 compartment, VA produced the largest cluster expansion observed, increasing the STAT1/B2M-defined CD4 N-3 cluster from 0.5% to 15.3% (29-fold), compared with 18-fold under BA and 11-fold under PA (**Supplementary Figures S3-21**). Within-cluster transcriptional changes under VA included upregulation of AP-1/IEG (JUN, FOS, JUNB, JUND) together with DUSP1 across 11 of 14 CD8 and 11 of 18 CD4 subclusters. A canonical IEG gene set was significantly enriched in VA-upregulated genes in every cluster tested (hypergeometric p=1e-4 to 1e-12). VA also induced MHC class II machinery transcripts (HLA-DRA, HLA-DRB1, HLA-DPA1, HLA-DPB1, HLA-DMA, CD74) across multiple subclusters.

BA drove a distinct resident memory phenotype, including the CD8 resident memory cluster (ZNF683⁺ with high cytotoxic granule expression), which expanded 10-fold under BA, compared with 4-fold under VA and 2.7-fold under PA. BA was also the strongest inducer of ZNF683 mRNA across CD8 subclusters. PA produced the most pronounced commitment to lowMT states. CD8 EM/CTL-lowMT expanded 9-fold, while multiple CD4 lowMT clusters (N-LowMT, Tfh-LowMT, EM-lowMT) increased 5.8-to 8.5-fold. Overall, 20% of CD4 and 5% of CD8 cells under PA occupied lowMT clusters (versus 3% and 0.6% under AXI). The median per-cell mitochondrial gene fraction was reduced under both PA and VA relative to AXI and BA.

A complete granular characterization of these shifts, including baseline cluster-defining differentially expressed genes for every CD4⁺ and CD8⁺ subcluster, within-cluster VA-vs-AXI differential expression in each subcluster, and Blood Transcription Module and Gene Ontology pathway enrichment per subcluster, is provided in **Supplementary Figures S3-21** and **Supplementary Table 3**. At equimolar 1 mM ex vivo dosing, the three SCFAs produced cellular phenotypes that were both functionally overlapping (broad improvement of cytotoxic, MHC class II, and memory-associated programs across most subclusters) and individually distinct: VA redistributed cells toward activated-naïve and memory states (CD4 N-3, CD8 N-1, CD8 CM-2, CD8 resting memory) and induced a broad AP-1/IEG response paired with DUSP1; BA committed cells preferentially toward resident memory differentiation; and PA distinctively committed multiple cell types to lowMT cell-state variants. We next examined the transcription factor and chromatin basis of these effects.

### Three SCFAs are associated with distinct patterns of chromatin accessibility and transcription factor activity in CAR T-cells

To investigate chromatin-level changes associated with SCFA conditioning, bulk ATAC-seq was performed on CAR T-cell products expanded in the presence of VA, BA, PA, or vehicle control (AXI; two biological replicates per condition from a single healthy donor). Differential accessibility was assessed using edgeR, transcription factor footprinting was performed using TOBIAS^30^ with JASPAR2024 motifs, and motif enrichment was evaluated using HOMER^31^.

At a stringent peak-level threshold, each SCFA produced markedly distinct chromatin remodeling profiles compared to vehicle (**Figure 4A**). VA produced a small but strongly opening**-**biased response (325/350 differentially accessible peaks opened; 93% directional). BA produced the largest overall chromatin remodeling but was symmetric to closing-biased (2,037 opened/2,817 closed; 42% opened). PA produced an intermediate-magnitude, mildly closing-biased response (383/497; 44% opened). The predominantly opening-biased pattern observed with VA was not seen under BA or PA.

**Figure 4.**
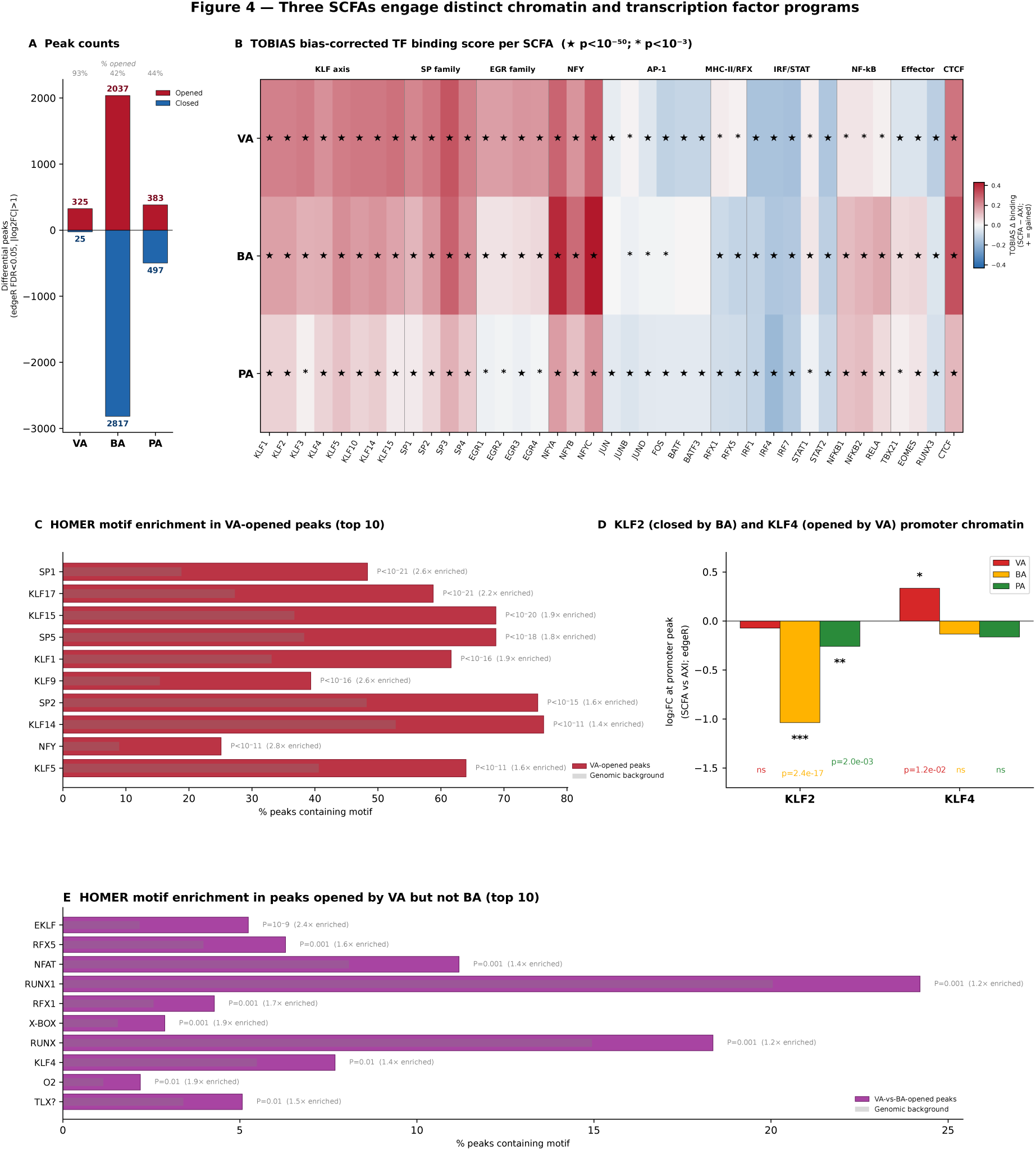
Valerate, butyrate, and propionate engage distinct chromatin and transcription factor programs. TOBIAS was run on per-condition merged-replicate ATAC-seq BAM files against the JASPAR2022 vertebrate motif library to quantify per-transcription-factor differential occupancy in each SCFA condition versus vehicle (U0/AXI). **(A)** Opened and closed peak counts of CAR T-cells treated with VA, BA, or PA. **(B)** Heatmap of TOBIAS bias-corrected transcription factor binding scores for 39 transcription factors (rows, color-coded by family) across VA, BA, and PA. Asterisks indicate TOBIAS-highlighted hits (p << 10⁻¹⁶⁰). EGR family categorical specificity: EGR1, EGR3, and EGR4 reach TOBIAS-highlighted significance (★) only under valerate; corresponding butyrate and propionate bars are sub-threshold despite measurable occupancy gains. **(C)** HOMER motif enrichment in VA-opened peaks. **(D)** Divergent KLF axis at promoter peaks: KLF4 chromatin opens preferentially under VA while KLF2 chromatin closes preferentially under BA. **(E)** HOMER motif enrichment in peaks opened by VA but not BA.

TOBIAS bias-corrected footprinting across a panel of 39 T-cell–relevant transcription factors indicated differences in inferred transcription factor activity between the SCFAs (**Figure 4B**). VA was associated with gains in binding at multiple KLF, SP, and EGR family motifs as well as NFY motifs. BA engaged a partially overlapping but distinct program centered on NFY and CTCF family motifs and smaller gains at TBX21 and EOMES that were not observed under VA. PA produced more limited shifts, primarily within the KLF/SP/EGR families. Reduced binding at IRF and AP-1 family motifs were observed across all three SCFAs, with the smallest reduction seen under VA.

HOMER motif enrichment in condition-specific peak sets was broadly consistent with the TOBIAS results (**Figure 4C**). VA-opened peaks showed strong enrichment for SP and KLF family motifs, whereas BA-opened peaks were enriched for NF-κB and other motifs, and PA-opened peaks showed enrichment for NFY motifs.

The single locus that most distinguished VA from BA was the KLF2 promoter^32^ (**Figure 4D**). At individual loci, BA conditioning was associated with reduced accessibility at the KLF2 promoter (log₂FC=−1.04, padj=2.5×10⁻¹⁷ vs AXI), while VA showed no significant change at that site. Only VA was associated with increased accessibility at the KLF4 promoter. Pairwise comparisons of VA-versus BA-opened peaks identified motifs enriched only under VA, including several KLF family members as well as RFX^33^ (X-box binding transcription factors central to MHC class II expression) and NFAT motifs (**Figure 4E**).

Together, TOBIAS footprinting and HOMER motif enrichment converged on a coherent set of transcription factor programs distinguishing the three SCFAs. VA engages the KLF/SP/EGR family at GC-rich promoter elements (asymmetric opening + KLF/SP motif enrichment + TOBIAS footprint gain) with a VA-distinguishing set of RFX, NFAT, and KLF1/4 motifs in VA-but-not-BA-opened peaks. BA produces a larger overall chromatin response, with KLF2 promoter closure consistent with effector commitment and NF-κB motif enrichment. PA produces an intermediate program centered on NFY motifs without effector commitment signatures. Taken together, these analyses identify three distinct patterns of chromatin accessibility and inferred transcription factor activity associated with the three SCFAs.

### Valerate is metabolized by CAR T-cells and restores anti-tumor function in vivo

In our previous study, we found that SCFA supplementation of CAR T-cell culture media improved tumor-cell killing under repeated antigen re-exposure^10^. To determine whether the chromatin and transcription factor changes documented above reflect direct valerate uptake and metabolism rather than indirect or off-target effects, we performed untargeted polar metabolomics on CD4-sorted and CD8-sorted CAR T-cell products from three independent healthy donors per condition (**Figure 5A**). VA-conditioned cells showed clear evidence of valerate uptake and mitochondrial β-oxidation: short-chain acylcarnitines (C4, C6, propionylcarnitine) and L-carnitine were elevated 2.0–3.4-fold in both the CD4 (p=0.045) and CD8 (p=0.012) compartments. Acetylcarnitine, the carnitine ester of mitochondrial acetyl-CoA, was elevated 2.83-fold in CD4 (p=0.10) and 2.63-fold in CD8 T-cells (p=0.005), consistent with valerate-derived carbon entering the mitochondrial acetyl-CoA pool. NAD⁺ was concurrently reduced 0.63-fold in CD4 (p=0.014) with the same directional change in CD8, consistent with VA-derived acetyl-CoA fueling histone acetyltransferase activity while NAD⁺-dependent sirtuin (class III HDAC) consumption is accelerated. Effector-associated metabolites including creatine, glutathione, spermidine, phosphocholine, aspartate, and L-proline were broadly elevated in VA-conditioned CAR T-cells (**Figure 5B**). These observations independently confirm in human CAR T-cells the dual epigenetic–substrate mechanism recently described for valerate in murine CAR T-cell models^34^, in which valerate acts simultaneously as a class I/II HDAC inhibitor, a dual acetyl-CoA/propionyl-CoA TCA-cycle substrate, and an NAD⁺/sirtuin substrate.

**Figure 5.**
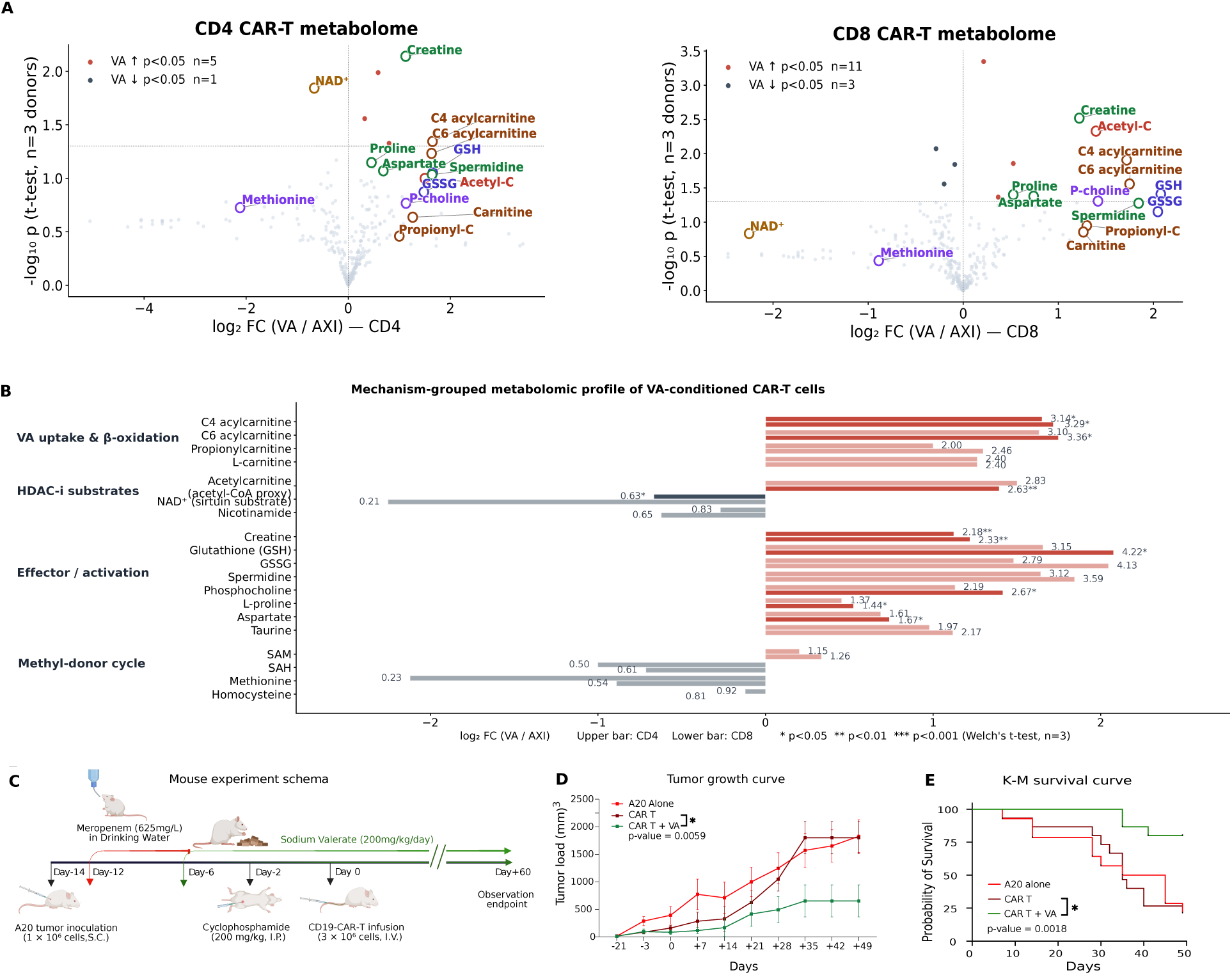
Valerate conditioning engages an intracellular metabolic program in CAR-T cells, and dietary valerate rescues CAR-T antitumor function in microbiome-depleted hosts in vivo. **(A)** Untargeted metabolomic profile of valerate-conditioned CAR-T cells. Volcano plots of VA-vs-AXI metabolite changes in sorted CD4⁺ (left, 5 up / 1 down at p < 0.05) and CD8⁺ (right, 11 up / 3 down) CAR-T cells; n = 3 donors per condition, paired t-test. Labels are color-coded by class: acylcarnitines (brown: C4, C6, propionylcarnitine, acetylcarnitine, carnitine — the dual TCA-entry signature of valerate β-oxidation), effector/biosynthetic metabolites (green: creatine, proline, aspartate, spermidine), redox glutathione axis (blue: GSH, GSSG), NAD⁺ / sirtuin substrate (yellow), and methyl-donor cycle (purple: methionine, phosphocholine). Creatine and acylcarnitines are the top VA-elevated hits in both compartments; NAD⁺ and methionine are coordinately depleted, consistent with valerate functioning as both a TCA-cycle substrate and a class I/II HDAC inhibitor that accelerates NAD⁺ and SAM consumption. **(B)** Mechanism-grouped intracellular metabolomic profile of VA-conditioned versus AXI CAR-T cells across three donors, with CD4 (upper bars) and CD8 (lower bars) compartments shown for each metabolite. Metabolites are grouped by mechanism: VA uptake and β-oxidation (acylcarnitines and L-carnitine), HDAC-inhibitor substrates (acetylcarnitine, NAD⁺, nicotinamide), effector and activation metabolites (creatine, glutathione, spermidine, phosphocholine, amino acids), and methyl-donor cycle (SAM, SAH, methionine, homocysteine). Asterisks indicate Welch’s t-test significance. **(C)** Mouse experiment schema: BALB/c mice received subcutaneous A20 lymphoma on day −14, oral meropenem (625 mg/L drinking water) from day −12 through day −6, dietary sodium valerate (200 mg/kg/day) from day −6 onward, cyclophosphamide preconditioning on day −2, and CD19-CAR-T infusion on day 0. **(D)** Tumor growth curves across the three arms (A20 alone, CAR-T, CAR-T + dietary valerate), with dietary valerate significantly reducing tumor burden over CAR-T alone (p = 0.0059). **(E)** Kaplan-Meier overall survival across the three arms, with dietary valerate significantly extending survival over CAR-T alone (log-rank p = 0.0018).

We next tested whether dietary supplementation of valerate could restore CAR T-cell anti-tumor function in vivo in a dysbiotic host. We previously demonstrated in a syngeneic A20 BALB/c lymphoma mouse model that oral meropenem exposure abolishes anti-CD19 CAR T-cell anti-tumor activity and produces a systemic multi-metabolite-deficient state in mouse serum that recapitulates the dysbiotic patient^10^. To test whether dietary sodium valerate supplementation could restore CAR T-cell efficacy in this model, we compared meropenem-treated A20 tumor-bearing mice that received no treatment, CAR T-cells alone, or CAR T-cells and dietary sodium valerate (**Figure 5C**). Mice receiving dietary valerate combined with CAR T-cells exhibited significantly reduced tumor burden relative to those receiving CAR T-cells alone (p=0.0059; **Figure 5D**) and extended overall survival (log-rank p=0.0018; **Figure 5E**), with valerate-treated animals maintaining tumor control through the day 60 observation endpoint.

## Discussion

The cellular composition and transcriptional state of the CD19-CAR T-cell product are major determinants of clinical response in LBCL^18,35–37^. Here, we show that a dysbiotic gut microbiome imprints a functionally impaired cellular state on the CAR T-cell apheresis product. However, replenishing a single depleted microbial metabolite, valeric acid, either during ex vivo manufacturing or through dietary supplementation can partially rescue CAR T-cell product quality and enhance antitumor activity. These results establish microbial metabolites as a genuine, actionable lever for improving CAR T-cell therapy and demonstrate that they are not merely correlative biomarkers but can be harnessed therapeutically.

This work makes three principal contributions. First, low stool valerate indicates a dysbiotic gut state that imprints a CD4-skewed CAR T-cell infusion product with diminished effector function. When these cells are subsequently infused into patients with pre-existing gut dysbiosis and chronic inflammation, their already compromised cytotoxic capacity may be further limited, contributing to inferior tumor control.

Second, valerate, butyrate, and propionate were each associated with distinct patterns of chromatin accessibility and transcription factor activity that align with the various cellular phenotypes observed by scRNA-seq. Valerate conditioning produced a predominantly opening-biased chromatin response and was linked to gains in KLF, SP, EGR, and NFY family transcription factor activity. These changes coincided with broad induction of AP-1/IEG transcripts and MHC class II machinery genes across multiple CAR T-cell subclusters. However, genome-wide AP-1 motif occupancy was reduced under all three SCFAs, indicating that the relationship between AP-1 transcript induction and actual transcription factor binding is complex and not fully resolved by bulk chromatin profiling. Butyrate, in contrast, drove substantially broader chromatin remodeling and was associated with gains in TBX21 and EOMES activity together with closure of the KLF2 promoter. This chromatin event has previously been linked to the release of CD8 T-cells from lymph node recirculation and progression toward effector states^38,39^. Propionate produced a more limited chromatin signature characterized by NFY motif enrichment and was distinguished by preferential commitment of both CD4 and CD8 T-cells to low-mitochondrial-content states, accompanied by a measurable reduction in per-cell mitochondrial gene expression. These findings suggest that each SCFA engages partially non-overlapping chromatin and transcription factor programs that may contribute to their distinct effects on CAR T-cell composition and transcriptional states. However, because these analyses were performed on bulk populations, the extent to which these chromatin-level changes occur within specific cell subclusters showing phenotypic shifts remains to be determined. Further studies will be needed to establish causal links between the observed chromatin programs and the functional consequences of SCFA conditioning.

Third, the stemness-associated changes observed under valerate were accompanied by opening at the KLF4 promoter, rather than by activation of the canonical TCF7/LEF1 axis. In fact, TCF7 motif occupancy appeared modestly reduced under valerate. These observations suggest that valerate may influence the retention of differentiation reserve through KLF4-related mechanisms, as opposed to the classical Wnt/TCF7 pathway typically associated with T-cell stemness. However, whether KLF4 directly contributes to any stemness-preserving effects of valerate cannot be concluded from chromatin profiling alone. Targeted functional experiments, such as KLF4 perturbation, are needed to establish its biological relevance.

A central methodological consideration is that all three SCFAs were tested at equimolar 1 mM concentrations. Because longer-chain SCFAs are more potent class I/II HDAC inhibitors at equimolar doses, the graded VA > BA > PA pattern observed for KLF, SP, and NFY family occupancy could reflect either SCFA-specific mechanisms or a simple difference in HDAC-inhibitory potency. Conclusions that remain robust to this caveat include direction-specific effects (KLF2 promoter closure by butyrate but not valerate), threshold effects unique to one SCFA (TOBIAS-significant TBX21/EOMES/NF-κB engagement under butyrate alone), and cell-state commitment not predicted by HDAC potency (the propionate-specific low-mitochondrial-content phenotype). Disentangling dose-equivalency from SCFA-categorical effects will require further HDAC-equipotent dose–response studies with acH3K27 quantification and controls using canonical HDAC inhibitors.

Overall, our findings indicate stool valerate as a viable bedside biomarker in which low stool valerate at CAR T-cell eligibility may identify patients whose infusion products carry a dysbiosis-imprinted signature. Ex vivo valerate conditioning of healthy-donor CAR T-cells produced a distinctive chromatin and cellular phenotype consistent with productive effector function. Whether ex vivo valerate can reverse the PIM-imprinted state in patient-derived material remains to be tested. In a meropenem-disrupted A20 lymphoma model, dietary valerate supplementation before apheresis reduced tumor burden and extended survival compared with CAR T-cells alone, indicating that pre-apheresis dietary intervention can improve CAR T-cell function in a dysbiotic host.

Our study has several limitations. The PIM apheresis analyses were observational and based on antibiotic exposure rather than randomized microbiome state, and the modest size of the PIM-exposed cohort (n=8 of 42 patients) limits power for individual transcriptional axes. ATAC-seq was performed on bulk populations, so it remains unknown whether the observed chromatin changes occurred within the same cellular subpopulations showing compositional or transcriptional shifts in the scRNA-seq analysis. We therefore cannot resolve the extent to which the chromatin-level differences functionally mediate cellular phenotypes. Cluster-resolved chromatin assays, including single-cell ATAC-seq or sorted-population ATAC, will be required to establish a direct mechanistic link between specific SCFA-driven transcription factor programs and corresponding T-cell states. Furthermore, only valerate was tested in vivo, while the in vivo effects of butyrate and propionate were not compared. All chromatin and transcriptional analyses were performed in healthy-donor CAR T-cells, and confirmation in patient-derived, particularly PIM-exposed, CAR T-cell products will be important for translation. Prospective randomized trials will ultimately be required to validate stool valerate as a biomarker and to establish the clinical benefit of metabolite-based interventions.

Overall, this work positions microbiome-derived metabolites at the interface of microbial ecology and cellular immunotherapy. The dysbiosis-imprinted signature in apheresis products and the distinct chromatin program engaged by ex vivo valerate conditioning identify mechanistic levers that may be relevant to other adoptive cell therapies. Prospective use of stool valerate as a companion diagnostic, combined with manufacturing-stage or dietary valerate interventions, offers a potential strategy for optimizing CAR T-cell therapy in patients with disrupted gut microbiomes.

## Declarations

### Ethics approval and consent to participate

All study procedures were performed in accordance with the Declaration of Helsinki and applicable institutional and federal guidelines. Plasma collection and clinical data abstraction at The University of Texas MD Anderson Cancer Center were performed under MD Anderson IRB-approved protocols. Written informed consent was obtained from all participants prior to specimen collection.

### Consent for publication

Not applicable. This article does not contain identifiable individual-level data, images, or videos.

### Authors’ contributions

N.Y.S. conceived and designed the study. L.R., S.G.R, A.R., K.S. has performed the *in vivo* validation studies. W.S. performed single-cell RNA sequencing, ATAC-seq data analysis, and transcription-factor regulon analysis. N.Y.S. led the bioinformatics and computational analysis of clinical microbiome cohort data, HUMAnN3 functional pathway analysis, and scRNA-seq analysis of apheresis products, and contributed to manuscript writing and figure preparation. F.D. performed all statistical analyses. J.F.F. conducted untargeted intracellular metabolomics profiling and analysis. A.J. performed ATAC-seq library preparation at the MD Anderson Epigenomics Profiling Core. C.K.S.T. provided German validation cohort patient specimens, clinical data, and metadata, and contributed to study concept and design. M.D.J. and R.R.J. contributed clinical specimens, data, and study design input from Moffitt Cancer Center and City of Hope, respectively. J.W., J.L., X.C., Y.T., S.P., M.D.R., F.C., J.C., C.L. and F.M.A. supervised patient recruitment and clinical data and biospecimen collection at MD Anderson. N.Y.S. and S.S.N. provided clinical oversight, resources, and contributed to study concept and design. M.R.T. edited the manuscript for language and readability. N.S., Y.Z., and S.S.N. jointly supervised this work, acquired funding, and critically reviewed and edited the manuscript. N.Y.S. drafted the manuscript. All authors critically reviewed and approved the final version of the manuscript.

## Acknowledgements

The authors thank the patients and their families for participating in this study, and the clinical and laboratory teams at The University of Texas MD Anderson Cancer Center, Moffitt Cancer Center and University Clinic Tuebingen, German Cancer Research Center, Germany for their support of specimen collection, processing, and clinical data abstraction.

