## Supplementary File for "Microbial valerate is associated with CAR T dysbiosis and its supplementation enhances CAR T function in B-cell lymphoma"

**Supplementary Methods**

### RESOURCE AVAILABILITY

Lead contact

Further information and requests for resources and reagents should be directed to and will be fulfilled by the lead contact, Neeraj Saini,. Materials transfer requires a completed Material Transfer Agreement (MTA) from The University of Texas MD Anderson Cancer Center (MDACC), Houston, USA.

Materials availability

The CD19-CAR-T construct (axicabtagene-ciloleucel-equivalent CD19-CD28-CD3z second-generation CAR; CD8-leader / CD8-hinge configuration) used in this study is the same product previously published by our group[1]; plasmid maps, viral preparation protocols, and product release criteria are described in that work and applied identically here unless noted. Sodium valerate (free salt) was obtained from Sigma-Aldrich (catalog 240370).

Data and code availability

Bulk ATAC-seq/ Single-cell RNA-seq raw and processed data are available from authors as per request. Untargeted metabolomics raw and processed data have been deposited earlier with a description given in the previous study[1]. Clinical microbiome shotgun-metagenomics taxon abundance tables and stool short chain fatty acid (SCFA) tables, with de-identified clinical metadata and per-patient survival outcomes, are provided in previous publications[2]. All custom analysis code and any additional information required to reanalyze the data reported in the current study is available from the lead contact upon request.

### EXPERIMENTAL MODEL AND STUDY PARTICIPANT DETAILS

Mouse in vivo CAR-T efficacy study

*Animals.* Female BALB/c mice aged 6 to 8 weeks were obtained from Jackson Laboratory and maintained under specific-pathogen-free conditions. All animal procedures were performed in accordance with protocols approved by the Institutional Animal Care and Use Committee protocol number 00002475-RN00**.**
*Tumor cell line and inoculation.* The murine B-cell lymphoma cell line A20 (American Type Culture Collection, TIB-208) was maintained in RPMI 1640 medium (Thermo Fisher Scientific, Waltham, MA; Cat No. 11875093) supplemented with 10% fetal bovine serum (RαD Systems by Biotechne, Cat No. S12450H), 50 μM β-mercaptoethanol (Thermo Fisher Scientific; Cat No. 21985023), 100 U/mL penicillin, and 100 μg/mL streptomycin (Thermo Fisher Scientific; Cat No. 15140122) at 37 °C in 5% $CO_2$. On day −14 of the experimental timeline, mice received subcutaneous inoculation on the right flank with $1 \times 10^6$ A20 cells suspended in 100 μL sterile phosphate-buffered saline (PBS; Thermo Fisher Scientific; Cat No. 10010049).

*Microbiome depletion.* Beginning on day −12, mice received oral meropenem (625 mg/L; MDACC, Lot no. 39AA0021-1, 39AA0025-1) dissolved in autoclaved drinking water and provided ad libitum through day −6 (six days of continuous antibiotic exposure). Drinking water containing meropenem was replenished every 48 to 72 hours with freshly prepared solution to maintain stable drug exposure.

*Sodium valerate supplementation.* Beginning on day −6, mice in the CAR-T plus valerate arm received sodium valerate (Med chem express, HY-W007087) at 200 mg/kg/day in autoclaved drinking water for the remainder of the experiment. Sodium valerate solutions were prepared fresh and replenished every 48 to 72 hours. Mice in the tumor alone and CAR-T alone arms received untreated drinking water from day −6 onward.

*Lymphodepleting preconditioning.* On day −2, all mice received a single intraperitoneal injection of cyclophosphamide (200 mg/kg; Sandoz, Cat No. 2905923) reconstituted in sterile saline.

*Murine CD19 CAR-T cell generation.* The detailed methods are provided in our previous manuscript[3]. Briefly, murine CD19-specific CAR-T cells were generated using the same CD19-CD28-CD3ζ retroviral construct described previously[3]. Splenocytes from BALB/c donor mice were activated with anti-CD3/anti-CD28 Dynabeads (Mouse T-Activator; Thermo Fisher Scientific, Cat No. 11132D) at a 1:1 bead-to-cell ratio for 24 hours. Cells were transduced after 24 hours with retroviral supernatant in the presence of Vectofusin-1 (Miltenyi Biotec, Cat No. 130-111-163) as per the manufacturer's protocol and expanded in complete RPMI 1640 supplemented with 100 IU/mL recombinant murine IL-2 (PeproTech, Cat No. 212-12) for 3 days. Transduction efficiency of CAR-T cells was confirmed by measuring the percentage of GFP+ cells by flow cytometry.

*CAR-T infusion and treatment groups.* On day 0, three treatment arms were established (n = 15 mice per arm): (1) tumor alone, receiving no CAR T-cells and no valerate; (2) CAR T-cells alone without valerate; and (3) CAR T-cells plus sodium valerate, receiving CAR T-cell infusion with continuous dietary sodium valerate supplementation as described above. Mice in the CAR T-cell -treated arms received $3 \times 10^6$ CD19 CAR T-cells in 200 μL PBS by tail vein intravenous injection. CAR T-cell transduction efficiency was confirmed at 60% by flow cytometry prior to infusion.

*Tumor monitoring and humane endpoints.* Tumor dimensions were measured every two to three days by digital caliper, and tumor volume was calculated as $V = (\text{length} \times \text{width}^2) / 2$. Body weights were monitored at each measurement. Mice were euthanized when tumors reached the institutional humane endpoint of 2,000 $mm^3$, when body weight loss exceeded 20%, or when other humane endpoint criteria were met (tumor ulceration, hunched posture, lethargy, dehydration). Surviving mice were observed through day 60 post-CAR T-cell infusion.

*Statistical analysis.* Overall survival was estimated by the Kaplan-Meier method and compared between treatment arms by log-rank (Mantel-Cox) test. P-values less than 0.05 were considered statistically significant. Statistical analyses were performed in GraphPad Prism v10.

Healthy human donors (CAR-T manufacturing for ex vivo mechanism)

Healthy donor peripheral blood mononuclear cells were isolated by Ficoll-Paque PLUS (Cytiva, Cat# 17144002) density gradient centrifugation. T-cells were activated with the ImmunoCult Human CD3/CD28/CD2 T Cell Activator (Stemcell Technologies, Cat# 10970) according to the manufacturer's protocol in complete RPMI 1640 medium supplemented with 300 IU/mL recombinant human IL-2 (GenScirpt, Cat No. Z00368-10 ), and were transduced on day 2 of expansion with a second-generation CD19–CD28–CD3ζ CAR lentivirus. Transduction efficiency was confirmed at ≥ 30% by flow cytometry, and product release criteria were applied as previously published[11]. Cells were expanded for 7 days. SCFA conditioning began on day 1 post-transduction and continued throughout expansion at concentrations approximating the upper end of physiologic systemic exposure (Supplementary Table S1A): 1 mM butyric acid (BA) (Sigma-Aldrich , Cat No. B103500), 1 mM propionic acid (PA) (Sigma-Aldrich, Cat No. 402907), or 1 mM valeric acid (VA) (Sigma-Aldrich, Cat No. 240370), each delivered as the sodium salt. The vehicle control arm (AXI) received sodium-buffered medium without SCFA. Media and SCFA were refreshed every 48 to 72 hours at a fixed T-cell concentration. One CAR T-cell manufacturing run was performed per donor.

Bulk ATAC-seq

ATAC-seq libraries were prepared at the MD Anderson Epigenomics Profiling Core following the Omni-ATAC protocol, as previously described[4,5]. Briefly, 50,000 viable cells were lysed in cold lysis buffer (10 mM Tris-HCl pH 7.4, 10 mM NaCl (Sigma-Aldrich; Cat# S9888), 3 mM $MgCl_2$ (Sigma-Aldrich; Cat# M1028), 0.1% NP-40

(Sigma-Aldrich; Cat# I8896), 0.1% Tween-20 (Sigma-Aldrich; Cat# P9416), 0.01% digitonin (Sigma-Aldrich; Cat# D141)) and the resulting nuclei were incubated with Tagment DNA TDE1 enzyme (Illumina) for 30 minutes at 37 °C. Tagmented DNA was purified using the Qiagen MinElute Reaction Cleanup Kit, amplified by PCR with custom barcoded primers, and size-selected with Ampure XP beads. Libraries were quantified by Qubit and visualized on a Bioanalyzer or TapeStation and pooled at equimolar concentrations. Sequencing was performed on an Illumina NovaSeq 6000 with 2 × 100 bp paired-end reads to obtain at 50 million high quality mapping reads per samples. Two biological expansion replicates per SCFA arm (AXI / BA / PA / VA) were processed.

Reads were adapter- and quality-trimmed (Trim Galore, Q20), aligned to GRCh38 with Bowtie2 (--very-sensitive, -X 2000), and filtered to retain properly-paired, primary, MAPQ ≥30, non-mitochondrial, non-duplicate fragments (samtools, Picard MarkDuplicates). Tn5 cut-site offsets (+4/−5) were applied. Per-sample peaks were called with MACS2 (--nomodel --shift −75 --extsize 150, $q < 0.01$). A consensus peak set was generated by merging peaks present in ≥2 samples (bedtools merge). Reads were counted per consensus peak per sample (featureCounts), normalized using DESeq2 size factors, and tested for differential accessibility per SCFA-vs-AXI contrast with DESeq2 (Wald test, Benjamini–Hochberg FDR). A peak was called significant at $|\log_2 FC| \geq 1$ and FDR < 0.05; per-condition counts were re-derived against the v2-corrected sample sheet after BA↔PA wet-lab labeling was reconciled (VA 4,631 opened / 2,389 closed; BA 1,286 / 336; PA 11,671 / 11,843 vs AXI). Motif enrichment over differential peaks vs background-matched genomic regions used HOMER with the HOCOMOCO v11 vertebrate library and GC-content-matched backgrounds, reporting $\log_2$ enrichment and Benjamini–Hochberg-adjusted hypergeometric p-values.

ATAC quality control

Quality control of ATAC-seq libraries was performed following ENCODE recommendations. Raw reads were trimmed and aligned to GRCh38/hg38 with Bowtie2. Mitochondrial reads, duplicate reads, low-quality alignments (MAPQ < 30), and ENCODE blacklisted regions were removed. Library complexity was assessed by NRF (Non-Redundant Fraction), PBC1, and PBC2 metrics on aligned mate-pair counts. TSS enrichment was calculated on 5,000 canonical protein-coding TSS sites with a 2 kb background window. Fraction of reads in peaks (FRiP) was calculated on the consensus peak set. Per-sample QC metrics are provided in Supplementary Table S4.

All eight samples (two biological replicates per condition: U0, VA, BA, PA) passed the ENCODE acceptable thresholds for TSS enrichment (>7; observed range 35.92–41.83, all classified "IDEAL") and FRiP (>0.2; observed range 0.42–0.60). Mitochondrial read fraction was 1.68–2.66%; bowtie2 mapping rate was 86.05–87.58%; post-filter read retention was 37.90–44.45%. Library complexity was acceptable (NRF > 0.7, PBC1 > 0.7, PBC2 > 3) in seven of eight samples; one sample (VA_R2) had borderline values (NRF 0.629, PBC1 0.684) reflecting higher sequencing depth (112.2 million raw reads versus 55.1–79.5 million in other samples), with PBC2 (3.35) and TSS enrichment (37.71) within acceptable range. This sample was retained.

**TOBIAS analysis**

To resolve cell-state-independent transcription factor (TF) binding from the bulk chromatin signal we applied TOBIAS (v0.16), bias-corrected ATAC-seq foot printing, to AXI-, BA-, PA-, and VA-conditioned CAR T-cell samples. Per-condition BAM files (deduplicated, filtered, Tn5-shifted as above) and the consensus peak set were used as inputs. ATACorrect was used to learn and remove the empirical Tn5 sequence-insertion bias per sample, producing per-base bias-corrected cut-site signal. ScoreBigwig then computed footprint scores across all consensus peaks, and the resulting per-condition bigWigs were used as input to BINDetect, which scans every consensus peak for matches to the JASPAR2022 CORE vertebrate motif library and computes, for each TF and each pairwise condition contrast, a differential binding score (Δbinding = mean footprint score in condition A − mean footprint score in condition B at all motif occurrences in accessible chromatin) together with a permutationbased $-\log_{10}$(p-value). A TF was flagged as "highlighted" by TOBIAS if it exceeded the joint significance threshold (|Δbinding| > 0.10 and $-\log_{10}$ p > 10) in at least one SCFA-vs-AXI contrast — this internal flag was used to define a high-confidence SCFA-responsive TF set for downstream interpretation. Three contrasts were the focus of mechanistic analysis: BA-vs-AXI, PA-vs-AXI, and VA-vs-AXI. Differential binding tables were merged with HOCOMOCO/JASPAR TF family annotations (KLF/SP, EGR, AP-1/bZIP, NFY, RFX, STAT, NK / homeobox) and visualized as (i) family-stratified Δbinding heatmaps across all three SCFAs, (ii) categorical bars marking which condition uniquely highlighted each TF, (iii) a VA-vs-BA scatter of Δbinding signed by direction (KLF4/KLF2 divergent axis), and (iv) AP-1 family cis-vs-trans deconvolution panels that separate motif-anchored AP-1 footprint changes (canonical AP-1 enhancer activity) from promoter-region GC-rich opening at AP-1-regulated loci (SP / KLF / EGR / NFY motifs falling within AP-1-target promoters), which is the dominant valerate behavior. Statistical comparisons across families were performed at the per-TF Δbinding level with Mann–Whitney U tests and at the per-sample peak level with paired Wilcoxon tests; multiple testing was controlled with Benjamini–Hochberg FDR. EGR1/EGR3/EGR4 reached TOBIAS-highlighted significance only in the VA contrast (categorically VA-specific), KLF4 was opened only by VA while KLF2 was closed only by BA (divergent KLF axis), and SP1/SP2 promoter motifs were uniquely opened by VA at AP-1 target-gene promoters. All scripts, parameter files, and BINDetect outputs are deposited with the manuscript.

Single-cell RNA-seq (scRNA-seq)

Three technical sequencing-library replicates per SCFA condition were generated from the donor HD-01 manufacturing run. Cells were partitioned and barcoded on the 10x Genomics Chromium platform (Single Cell 3' v3.1 chemistry). Libraries were prepared per the manufacturer's protocol and sequenced on Illumina NovaSeq 6000 to a target depth of >= 30,000 reads per cell. Reads were processed with Cell Ranger v7.2.0 against GRCh38-2020-A. Filtered cell-by-gene matrices were imported into Scanpy v1.9.6. Quality-control thresholds excluded cells with < 400 or >6,000 detected genes, or > 5% mitochondrial reads. After QC, ~10,000 cells per condition were retained (39,739 cells in total for AXI, BA, PA, and VA conditions).

Per-library normalization (Scanpy `normalize_total` to 10,000 counts followed by log1p transformation), highly-variable-gene selection (top 5,000 HVGs), and PCA (top 30 PCs) were performed independently per library. Batch effects are not observed in the integrated data across four conditions. Cluster annotation was performed on the basis of canonical $CD4^+$ and $CD8^+$ T cell lineage markers (CD4 for CD4 lineage; CD8A for CD8 lineage; CD27, SELL, TCF7, and CCR7 for naive; CXCR3, IL2RB, and ITGAL for central memory, KLRB1, ITGB1, CD38, and CCR5 for effector memory, GZMA, GZMB, PRF1, NKG7, GNLY, and CCL5 for cytotoxic; CD40LG, ICOS, and BCL6 for follicular helper; TBX21, IFNG, PRF1, and NKG7 for Th1, IL4, IL5, IL13, GATA3 and PTGDR2 for Th2; FOXP3, IKZF2, IL2RA, and TNFRSF18 for regulatory in Supplementary Figs. 1-3) plus unsupervised top-marker gene expression (Supplementary Figs. 4-5). Leiden clustering with resolution 2.0 is used for subclustering within $CD4^+$ or $CD8^+$ CAR-T population Per-cluster differential expression was computed with the Wilcoxon rank-sum test with Benjamini-Hochberg correction for false discovery rate control (`rank_genes_groups` function from *scanpy* python module). Thresholds for calling significant differentially expressed genes were deemed by $|\log_2 FC|$ >= 2 and adjusted p < 0.01. Per-cell module scores for the ten functional T-cell programs (i.e., activation, antigen presentation, cytotoxicity, exhaustion, interferon response, Isag, memory, persistence, proliferation, regulatory programs.

Pre-ranked GSEA per cluster on the cluster-specific fold-change ranking was performed with GSEApy v1.1.3 against the same gene-set collections (Gene Ontology Biological Process, KEGG, and Blood Transcription Module) used for ATAC-seq GSEA, enabling the synchronized ATAC-up / RNA-up integration shown in Figure 4C.

Transcription-factor target regulon construction

Eighteen literature-anchored T-cell-relevant TF target gene sets were assembled from primary publications: AP-1 / NFAT family (FOS / FOSB / FRA1 in T-cell activation[6]; JUN direct T-cell targets[7]; AP-1 T-cell targets per Macian 2005[8,9] ; NFAT targets per Macian 2005[8]; BATF targets per Glasmacher 2012[10]; STAT1 IFN-gamma targets (Schroder 2004); IRF1 antigen-processing targets (canonica[11]l); CIITA / RFX-family MHC class II targets (Reith 2005[12]); MHC class I and II machinery (canonical)[13,14] ; RUNX3 CD8-effector targets (Cruz-Guilloty 2009[14]); T-bet (TBX21) Th1 targets (Hwang 2005, Szabo 2002[15]); BACH2 memory-regulator targets ( Roychoudhuri 2016[16]); TCF7 / WNT naive-stemness targets (Gattinoni 2011[17]); NF-kB / RELA targets (Hayden 2008[18]). Pre-ranked GSEA on the VA-vs-AXI gene-level ATAC-seq ranking against this regulon collection used GSEApy v1.1.3 with 2,000 permutations, weighted enrichment statistic, seed = 42, and gene-set size 5-500. Significance was assessed at FDR < 0.05; results are in Figure 5 and Supplementary Table S3.

Cellular metabolomics

Cellular metabolomics analyses were conducted as previously described[3].

QUANTIFICATION AND STATISTICAL ANALYSIS

**Figure 1 — Cross-cohort microbiome and HUMAnN3 functional pathway analysis**

Stool SCFA concentrations were $\log_{10}$-transformed using $\log_{10}$(value + 1) to handle zero counts. Bacterial species relative abundance was profiled by MetaPhlAn3 for the MDACC discovery cohort (n = 69) and for the German validation cohort (n = 54). Within each cohort, species tables were filtered to species present in at least 5 samples and $\log_{10}$-transformed using $\log_{10}$(value + $10^{-6}$). HUMAnN3 (v3.6) community-level pathway abundance tables were filtered to remove unmapped and unintegrated rows and $\log_{10}$-transformed using $\log_{10}$(value + $10^{-3}$). Within each cohort, two-sided Spearman rank correlations between each species or pathway and stool valerate concentration were computed, and p-values were adjusted across all tested features within that cohort using the Benjamini-Hochberg false discovery rate (BH-FDR). Cross-cohort replication of a species or pathway was defined a priori as reaching BH-FDR < 0.10 in both cohorts with concordant sign of the Spearman coefficient. In the species-level analysis (Figure 1A, 1B), species reaching nominal $p < 0.05$ against stool valerate in at least one cohort were retained for display, and concordant replication was assessed in the top MDACC valerate-associated species available in both species catalogs. In the HUMAnN3 pathway-level analysis (Figure 1C), 84 of 528 detected pathways reached BH-FDR < 0.10 in MDACC, of which 23 also reached BH-FDR < 0.10 in the German cohort with concordant direction. Tertile-based dose-response analysis (Supplementary Figure S1) stratified MDACC patients into Low, Mid, and High stool valerate tertiles (n ≈ 23 per tertile). For each of the 23 cross-cohort replicating pathways, mean $\log_{10}$ pathway abundance was computed within each tertile and z-scored across the three tertile means to yield a row-standardized gradient. A pathway was classified as monotonically increasing if mean abundance was strictly higher in each successive tertile (Low < Mid < High) and monotonically decreasing if strictly lower.

**Figure 2 — Apheresis CAR-T cellular state and prognostic Cox regression**

The discovery scRNA cohort comprised 42 large B-cell lymphoma (LBCL) patients treated with axicabtagene ciloleucel and profiled at apheresis (pre-manufacture), of whom 24 (5 PIM-exposed, 19 no-PIM; 133,405 single cells) had cluster-resolution annotation from Deng et al. (Nat Med 2020). Patients were stratified into 8 with PIM exposure (piperacillin–tazobactam, imipenem, or meropenem within 30 days before apheresis) and 34 with no prior antibiotic exposure. All 42 patients contributed to per-cell module and PFS analyses; CAR-transgene-positive proportion (35.9%) and cell yield were comparable between groups.

Cluster-level compositional shifts (Figure 2A) were quantified across all 15 Deng-annotated apheresis CAR-T clusters by computing the patient-level fraction of each cluster, taking $\log_2$(PIM mean / no-PIM mean), and testing each cluster by two-sided Mann–Whitney U test on patient fractions. Clusters were colored by Deng prognostic class (poor-prognosis, good-prognosis, other) using the published Deng et al. cluster-prognosis assignment; nominal patient-level p-values are displayed. Compartment-level CD4 / CD8 partitioning (Figure 2B) was performed on patient-aggregated CD4 and CD8 fractions and compared between PIM (n = 8) and no-PIM (n = 34) groups by the same two-sided Mann–Whitney U test; box plots show median, IQR, and individual patient points.

Per-patient transcription-module scoring (Figure 2C) used Scanpy v1.9 sc.tl.score_genes over published, pre-cited gene sets assembled from the primary literature (full citations are provided in the main manuscript) to address potential team-curation bias. Each module was scored per cell, then aggregated to a patient-level median (cell-intrinsic, orange) or to a patient-level whole-blood signature (patient-level, blue). $\text{Log}_2$(PIM mean / no-PIM mean) was plotted with two-sided Mann–Whitney U p-values on patient values; no module was pseudoreplicated at the cell level. Significant modules at $p < 0.05$ (denoted *) included AP-1/transphosphor activation (p = 0.043), Crawford 2014 (p = 0.034), and Wherry 2007 exhaustion (p = 0.011); MHC class II antigen presentation was suppressed under PIM (p = 0.028).

Progression-free survival (PFS) analysis (Figure 2D) was performed at the patient level using the lifelines Python package v0.27. The composite chronic-activation score was computed as AP-1/IEG + Wherry-exhaustion − naive-memory module scores and z-scored across the 42 patients. Patients were partitioned into Low, Mid, and High chronic-activation tertiles (n = 14 per tertile); a Kaplan–Meier log-rank test compared survival across tertiles, and a Cox proportional-hazards model regressed PFS on the continuous z-scored composite, yielding HR = 1.56 (95 % CI 1.12–2.41, Wald p = 0.012). A multivariable Cox model additionally adjusted for age (continuous), sex (male vs female), Ann Arbor stage (III/IV vs I/II), prior lines of therapy (continuous), DLBCL histology (yes vs no), and PIM exposure. Hazard ratios with 95% CIs and Wald-test p-values are reported in Supplementary Figure S3 and Supplementary Table S3. This analysis included 42 patients with 28 progression events at a median follow-up of 24 months. All p-values are two-sided; BH-FDR was applied to multi-feature tests where indicated.

**Figure 3 — $CD4^+$ and $CD8^+$ CAR-T scRNA under four-arm SCFA conditioning**

Healthy-donor CD19-CAR T-cell products were conditioned with vehicle (AXI), butyrate (BA), propionate (PA), or valerate (VA) at 1 mM for 7 days during the full manufacturing process and profiled by droplet-based single-cell RNA-seq (10x Genomics 3′ v3.1). FASTQs were aligned to GRCh38 with Cell Ranger v7.1 (CAR-transgene reference appended). Per-sample counts were imported into Scanpy v1.9 and concatenated. Cells were retained at 500 ≤ genes/cell ≤ 6,000 and percent-mitochondrial ≤ 15%; doublets were removed by Scrublet (threshold 0.25). Counts were normalized to $10^4$ per cell, log1p-transformed, and 3,000 highly variable genes were selected by Seurat-flavored HVG. The expression matrix was scaled (clip ± 10), PCA-reduced to 50 components, batch-corrected across donors and conditions with Harmony, and embedded by UMAP (n_neighbors = 30, min_dist = 0.3). $CD4^+$ and $CD8^+$ compartments were partitioned by CD4 / CD8A / CD8B expression and re-embedded independently. Sub-clustering used Leiden at resolution 2.0 ($CD4^+$, 18 subclusters) and 1.5 ($CD8^+$, 14 subclusters). Cluster identity was assigned from canonical lineage markers (naïve: TCF7, LEF1, SELL, CCR7, IL7R; central memory; effector memory; cytotoxic: GZMA/B, PRF1, NKG7, GNLY; Tfh, Th1, Th2, Treg; cycling: MKI67, histone, CCNB2) supported by dot-plot expression and module scoring (sc.tl.score_genes) of published canonical signatures. Cell-cycle phase (G1/S/G2M) was assigned by sc.tl.score_genes_cell_cycle. Low-mitochondrial sub-states (CD4 N-LowMT, Tfh-LowMT, EM-lowMT; CD8 EM/CTL-lowMT) were defined as clusters whose median mitochondrial module score fell > 2 MAD below the compartment median. Per-cluster baseline DEGs (AXI-only) and per-cluster VA-vs-AXI DEGs were computed with sc.tl.rank_genes_groups (Wilcoxon rank-sum, tie-corrected, BH-FDR across all genes); a gene was called significant at $|\log_2 FC| > 1$ (intra-naïve / intra-

CTL contrasts) or > 2 (cluster-vs-rest) and FDR < 0.01. Cluster-proportion shifts across SCFA arms were tested at the sample level (proportion of each cluster per sample) by two-sided Mann–Whitney U with BH-FDR across clusters within each contrast. Functional-axis radar scores (activation, antigen, cytotoxicity, exhaustion, interferon, Treg, memory, persistence, proliferation, regulatory) were computed per cell via sc.tl.score_genes using literature-anchored MSigDB Hallmark, Wherry exhaustion, and Gattinoni memory gene sets and aggregated to per-condition medians.

**Figure 4A-D Bulk ATAC-seq peak-level differential accessibility**

Peak counts and read assignments were generated as described in the Methods (peak calling, consensus peak set, fragment count matrix). Differential peak accessibility per SCFA versus vehicle (AXI) was computed using edgeR (v3.42) on the consensus peak count matrix. A negative-binomial generalized linear model with quasi-likelihood F-test (glmQLFTest) was applied with batch-aware design (~batch + condition); p-values were adjusted for multiple testing using the Benjamini-Hochberg false discovery rate (FDR). A peak was called differentially accessible at FDR < 0.05; for opening-direction summary statistics (Figure 4A) and locus-level reporting (Figure 4D), peaks were additionally required to have $|\log_2 FC| > 1$.

**Figure 4B TOBIAS bias-corrected foot printing**

Bias-corrected Tn5 foot printing was performed with TOBIAS (v0.16) against the JASPAR2024 CORE vertebrate motif library. ATACorrect was used for sequence-bias correction of merged per-condition BAM files; ScoreBigwig computed per-position footprint scores; and BINDetect computed per-motif differential binding scores for each SCFA-versus-AXI contrast. Motifs were called "TOBIAS-highlighted" at $p < 10^{-3}$ (asterisks in Figure 4B), with the most stringent significance threshold ($p < 10^{-50}$) marked separately (stars in Figure 4B). TOBIAS Δ binding score was inverted for visualization such that positive values indicate increased binding under the SCFA condition relative to AXI.

**Figure 4C and 4E HOMER motif enrichment**

Known-motif enrichment in differentially accessible peak sets was performed with HOMER (findMotifsGenome.pl, v4.11) against the human hg38 genome with the JASPAR2024 vertebrate motif library. The consensus peak set served as the GC-matched genomic background. For each contrast, opened peaks (edgeR FDR < 0.05, $\log_2 FC > 1$) were submitted as the target set. Motif enrichment was tested with HOMER's hypergeometric test; P-values are reported directly from the HOMER output ("P-value" column). Benjamini-Hochberg q-values were computed by HOMER and used for top-motif filtering (q < 0.05 unless otherwise stated). Fold enrichment was computed as (% target peaks with motif) / (% background peaks with motif).

**Figure 5A-B Metabolomics — volcano and mechanism-grouped statistics**

Untargeted polar metabolomics was performed on CD4-sorted and CD8-sorted CAR T-cell products from three independent healthy donors per arm (VA versus vehicle [AXI]). Raw peak intensities were $\log_2$-transformed and centered. For each metabolite, $\log_2$(VA/AXI) was computed per donor and tested by two-sided Welch's t-test (n = 3 donors per condition) without correction for multiple testing, given the targeted nature of the metabolite panel and the small number of biological replicates. Significance thresholds reported on bar charts: *p < 0.05; **p < 0.01; ***p < 0.001. Volcano plots (Figure 5A) display $\log_2$(VA/AXI) versus $-\log_{10}$(P) for each detected metabolite per compartment; metabolites passing p < 0.05 are highlighted.

**Figure 5D Mouse tumor growth curve**

Subcutaneous A20 lymphoma volume was measured by caliper through the observation endpoint or until predefined humane endpoint (tumor volume > 2,000 $mm^3$). Tumor volumes were calculated as length × $width^2$ × 0.5. Group sizes were as indicated in the figure legend. Group-wise comparison of tumor growth between the CAR T-cell–alone and CAR T-cell + dietary valerate arms was performed by two-way repeated-measures ANOVA on log-transformed tumor volume with arm and time as factors and animal as a random effect; $p = 0.0059$ for the CAR-T-alone versus CAR-T + VA contrast.

*Panels F-H: In vivo CAR-T efficacy.* Tumor volume was compared between arms by two-sided Mann-Whitney U test ($p = 0.0059$). Overall survival was estimated by Kaplan-Meier and compared by log-rank test ($p = 0.0018$). Both analyses were performed in GraphPad Prism v10.

Explained above

**Software summary**

ATAC-seq differential analysis was performed in R (v4.3) with edgeR (v3.42); TOBIAS footprinting in Python (v3.11) with TOBIAS (v0.16); HOMER motif enrichment with HOMER (v4.11); metabolomics statistics and visualization in Python (v3.11) with scipy.stats and pandas; mouse survival and tumor growth analyses in GraphPad Prism (v10) and R (v4.3) with the survival package (v3.5). MetaPhlAn3 v3.0.14 were used for taxonomic profiling and HUMAnN3 v3.6 for functional pathway profiling. Throughout the manuscript, asterisks denote: * $p < 0.05$, ** $p < 0.01$, *** $p < 0.001$ (specific test indicated in each panel). BH-FDR refers to the Benjamini-Hochberg false discovery rate within the indicated feature set.

KEY RESOURCES TABLE

Biological reagents and animal models

| Reagent or resource | Source | Identifier |
| --- | --- | --- |
| Anti-CD19 CAR retrovirus (CD19-CD28-CD3z, CD8-leader / CD8-hinge) | Gift from Marco Davila (Roswell Park Comprehensive Cancer Center) | |
| Anti-CD3 / anti-CD28 Dynabeads (mouse T-Activator) | Thermo Fisher | Cat# 11132D |
| Recombinant human IL-2 | Peprotech | Cat# 200-02 |
| A20 murine B-cell lymphoma cell line | ATCC | ATCC TIB-208 |
| Female Balb/c mice (6-8 weeks) | Jackson Laboratory | Strain code 028 |

Chemicals, peptides, and small molecules

| Reagent or resource | Source | Identifier |
|---|---|---|
| Sodium butyrate | Sigma-Aldrich | Cat# 303410 |
| Sodium propionate | Sigma-Aldrich | Cat# P1880 |
| Sodium valerate (free salt) | Sigma-Aldrich | Cat# 240370 |
| Meropenem | Sigma-Aldrich (or institutional pharmacy) | Cat# M2574 |
| Cyclophosphamide | Sigma-Aldrich (or institutional pharmacy) | Cat# C0768 |


1. Prasad, R., Rehman, A., Rehman, L., Darbaniyan, F., Blumenberg, V., Schubert, M.-L., Mor, U., Zamir, E., Schmidt, S., Hayase, T., et al. (2025). Antibiotic-induced loss of gut microbiome metabolic output correlates with clinical responses to CAR T-cell therapy. Blood *145*, 823-839. 10.1182/blood.2024025366.
2. Stein-Thoeringer, C.K., Saini, N.Y., Zamir, E., Blumenberg, V., Schubert, M.L., Mor, U., Fante, M.A., Schmidt, S., Hayase, E., Hayase, T., et al. (2023). A non-antibiotic-disrupted gut microbiome is associated with clinical responses to CD19-CAR-T cell cancer immunotherapy. Nat Med. 10.1038/s41591-023-02234-6.
3. Patel, R., Devashish, K., Singh, S., Nath, P., Gohel, D., Prasad, R., Singh, M., and Saini, N. (2021). Vectofusin-1–based T-cell transduction approach compared with RetroNectin-based transduction for generating murine chimeric antigen receptor T-cells. bioRxiv, 2021.2008.2028.458011. 10.1101/2021.08.28.458011.
4. Corces, M.R., Trevino, A.E., Hamilton, E.G., Greenside, P.G., Sinnott-Armstrong, N.A., Vesuna, S., Satpathy, A.T., Rubin, A.J., Montine, K.S., Wu, B., et al. (2017). An improved ATAC-seq protocol reduces background and enables interrogation of frozen tissues. Nat Methods *14*, 959-962. 10.1038/nmeth.4396.
5. Marin, D., Li, Y., Basar, R., Rafei, H., Daher, M., Dou, J., Mohanty, V., Dede, M., Nieto, Y., Uprety, N., et al. (2024). Safety, efficacy and determinants of response of allogeneic CD19-specific CAR-NK cells in CD19(+) B cell tumors: a phase 1/2 trial. Nat Med *30*, 772-784. 10.1038/s41591-023-02785-8.
6. Vahedi, G., Kanno, Y., Furumoto, Y., Jiang, K., Parker, S.C., Erdos, M.R., Davis, S.R., Roychoudhuri, R., Restifo, N.P., Gadina, M., et al. (2015). Super-enhancers delineate disease-associated regulatory nodes in T cells. Nature *520*, 558-562. 10.1038/nature14154.
7. Lynn, R.C., Weber, E.W., Sotillo, E., Gennert, D., Xu, P., Good, Z., Anbunathan, H., Lattin, J., Jones, R., Tieu, V., et al. (2019). c-Jun overexpression in CAR T cells induces exhaustion resistance. Nature *576*, 293-300. 10.1038/s41586-019-1805-z.
8. Macian, F. (2005). NFAT proteins: key regulators of T-cell development and function. Nat Rev Immunol *5*, 472-484. 10.1038/nri1632.

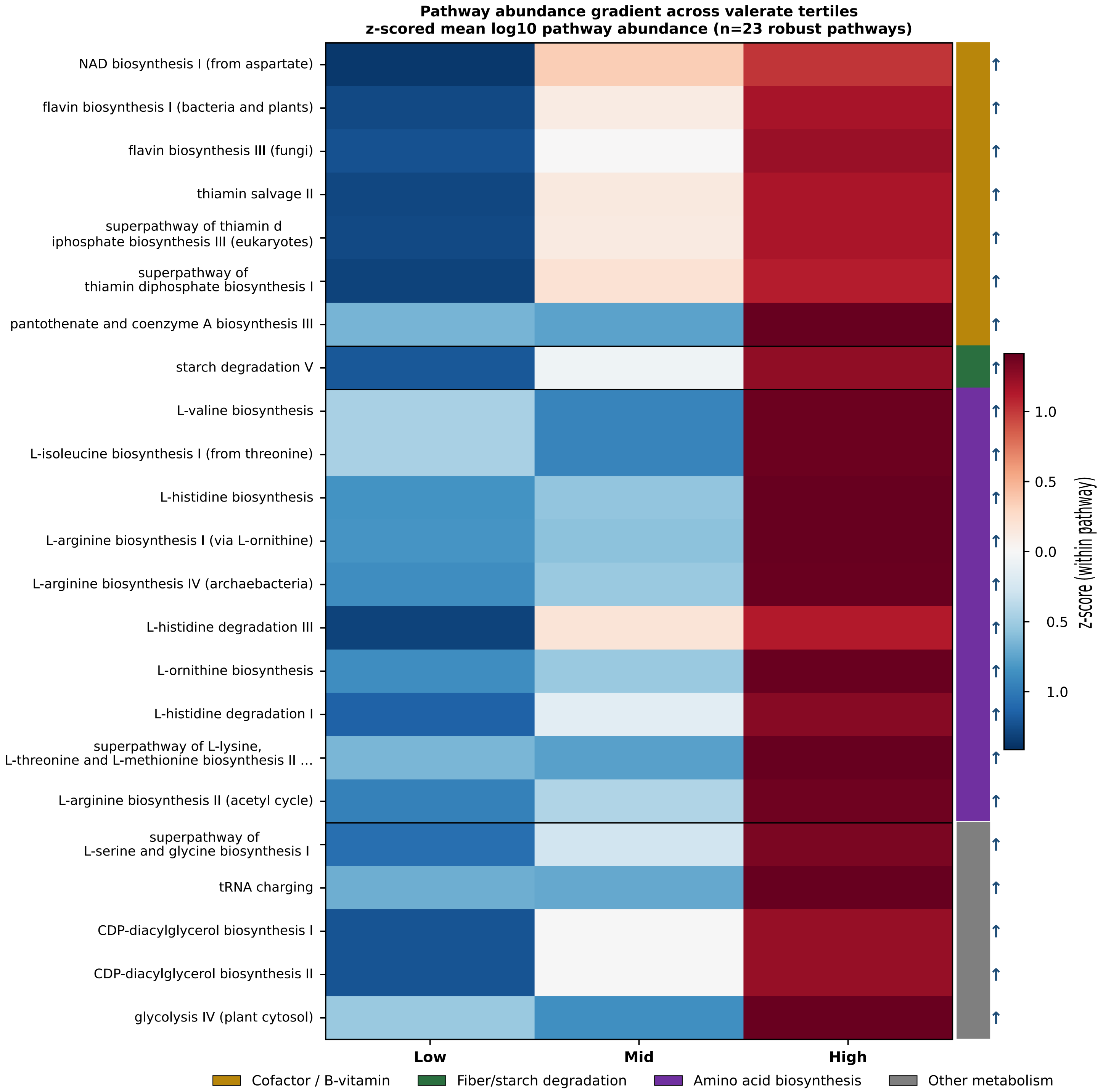


**Supplementary Figure S1.** Pathway abundance z-score across stool valerate tertiles in MDACC for the same 23 pathways shown in panel C. Cross-cohort microbiome functional pathway depletion across stool valerate tertiles defines a multi-metabolite-deficient gut state. Heatmap of z-scored mean $\log_{10}$ pathway abundance across stool valerate tertiles (Low, Mid, High) for the 23 HUMAnN3 functional pathways. Each row is a microbial functional pathway, each column is a valerate tertile, and color encodes within-pathway z-score. Seventeen of the 23 replicating pathways followed this monotonic dose-response, with no pathway showing the inverse direction.

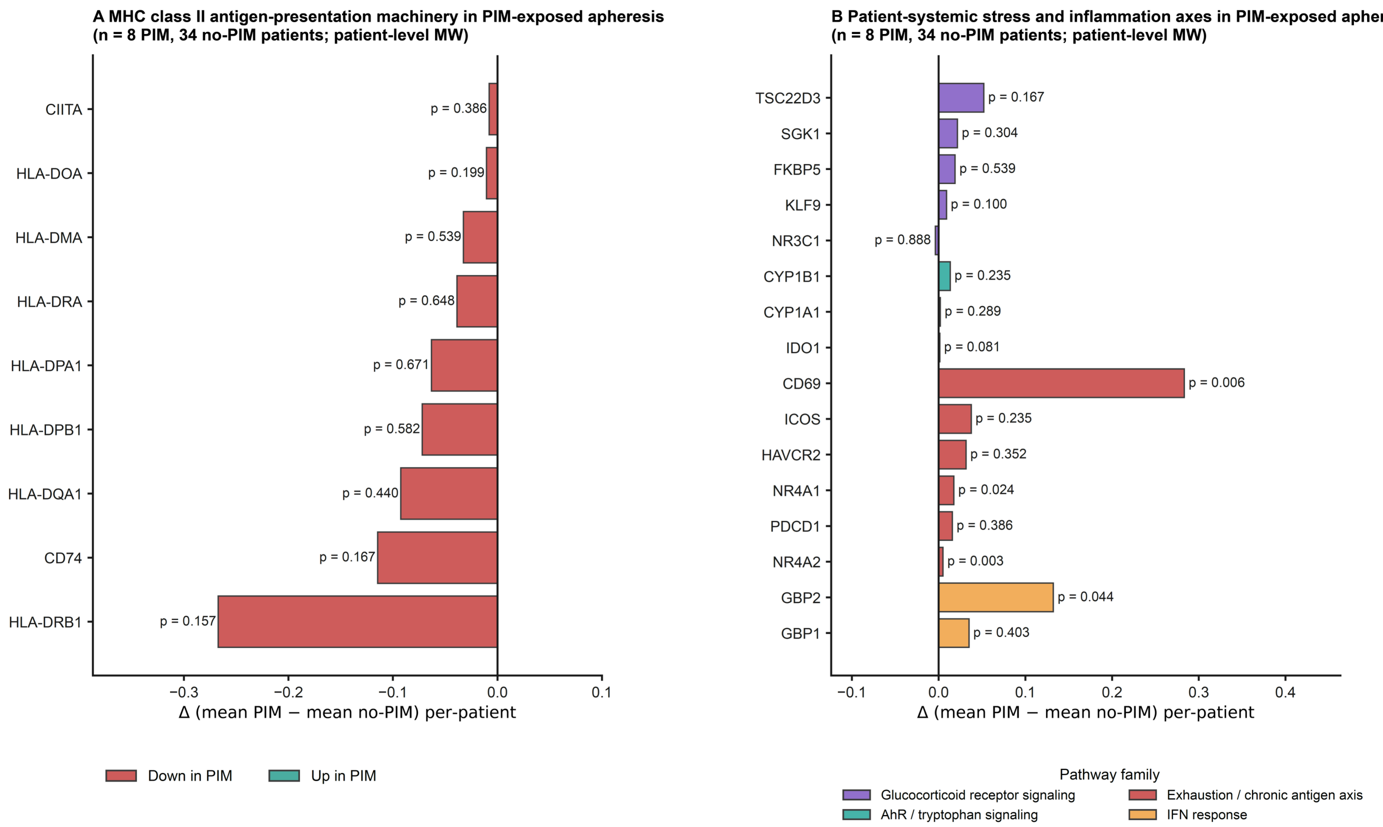


**Supplementary Figure 2. Gene-level patient-aggregated expression shifts in PIM-exposed apheresis at the constituent-gene resolution of the MHC class II and stress-axis signatures shown in Figure 2C.** For each gene plotted, per-cell normalized expression was extracted from the apheresis CAR-T product CD4 (n = 164,114 cells) and CD8 (n = 185,226 cells) compartments and aggregated to a per-patient mean across each patient's pooled CD4 and CD8 cells. Mann-Whitney U tests were performed at the patient level (n = 8 PIM-exposed and n = 34 unexposed patients), with raw two-sided p-values shown next to each bar. Patient-level Mann-Whitney is reported here rather than cell-level Mann-Whitney because the cell-level test treats correlated cells from the same patient as independent observations and produces inflated significance. **(A)** MHC class II antigen-presentation machinery genes in PIM-exposed apheresis. Bars show the difference in mean expression between PIM-exposed and unexposed patients (Δ = mean PIM − mean no-PIM) at the per-patient level for nine MHC class II machinery genes: HLA-DRB1, CD74, HLA-DQA1, HLA-DPB1, HLA-DPA1, HLA-DRA, HLA-DMA, HLA-DOA, and CIITA. All nine genes trend directionally lower in PIM-exposed apheresis (red bars, down in PIM), but no gene reaches per-patient raw $p < 0.05$ (smallest p = 0.157 for HLA-DRB1). **(B)** Patient-systemic stress and inflammation axes in PIM-exposed apheresis. Bars show the per-patient Δ for 16 genes spanning four patient-systemic pathway families, color-coded by family: glucocorticoid receptor signaling (FKBP5, NR3C1, TSC22D3, SGK1, KLF9, purple), aryl hydrocarbon receptor and tryptophan catabolism (CYP1A1, CYP1B1, IDO1, teal), T-cell exhaustion and chronic antigen receptor signaling (CD69, HAVCR2, ICOS, NR4A1, NR4A2, PDCD1, red), and interferon-γ response (GBP1, GBP2, orange). Four genes reach per-patient raw $p < 0.05$: NR4A2 (exhaustion, p = 0.003), CD69 (exhaustion, p = 0.006), NR4A1 (exhaustion, p = 0.024), and GBP2 (interferon response, p = 0.044). Three additional genes show borderline trends (IDO1 p = 0.081, KLF9 p = 0.100, TSC22D3 p = 0.167). No gene passes BH-FDR correction across the 25 simultaneously tested patient-level Mann-Whitney comparisons (lowest BH-FDR 0.078 for NR4A2 and 0.080 for CD69).

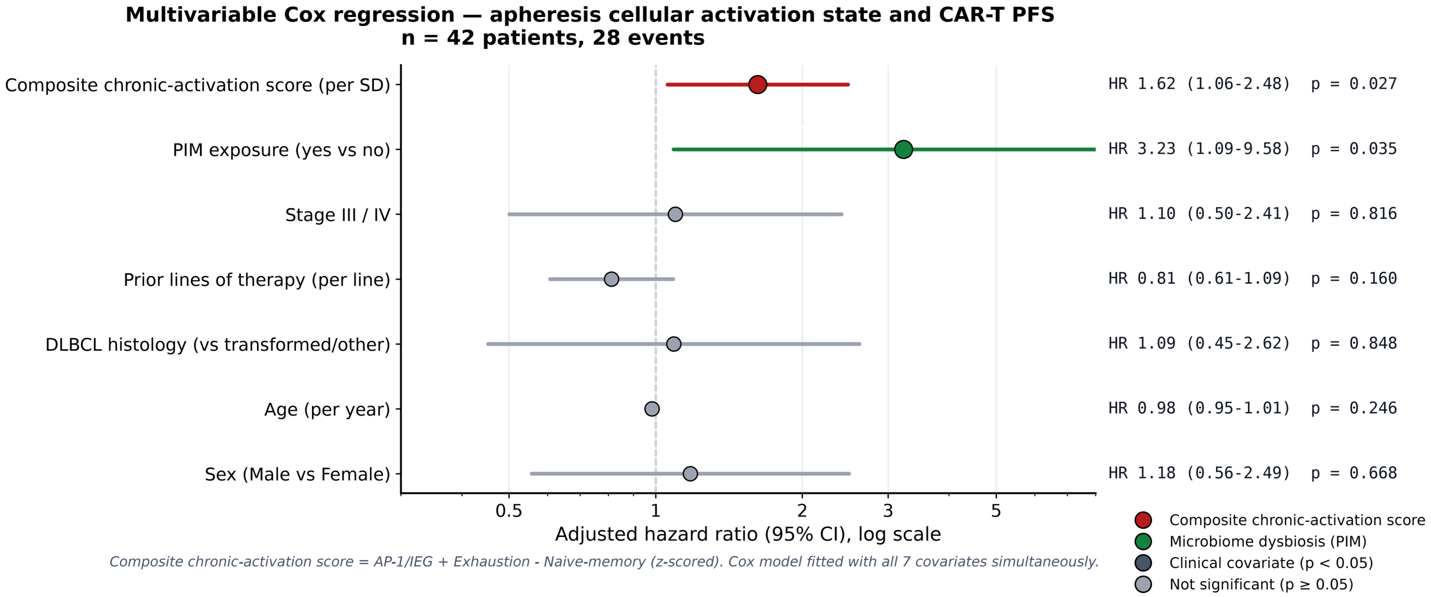


**Supplementary Figure S3.** Multivariable Cox regression confirms the apheresis cellular activation state as an independent predictor of progression-free survival after CAR-T therapy. The model includes all seven covariates simultaneously: the composite chronic-activation score (per standard deviation), PIM exposure (piperacillin-tazobactam, imipenem, or meropenem within 30 days of leukapheresis, yes vs no), Stage III/IV disease (vs I/II), prior lines of therapy (per additional line), DLBCL histology (vs transformed or other), age (per year), and sex (male vs female). The composite chronic-activation score is computed at the per-sample level as the sum of the AP-1 / immediate-early-gene module and the exhaustion module minus the naive-memory module, z-scored across the cohort. Adjusted hazard ratios are shown on a logarithmic x-axis with 95% confidence intervals, with the numeric estimate and p-value listed for each covariate. Both the composite chronic-activation score (HR 1.62, 95% CI 1.06 to 2.48, p = 0.027) and PIM exposure (HR 3.23, 95% CI 1.09 to 9.58, p = 0.035) remain independently associated with inferior progression-free survival after mutual adjustment for clinical risk factors.

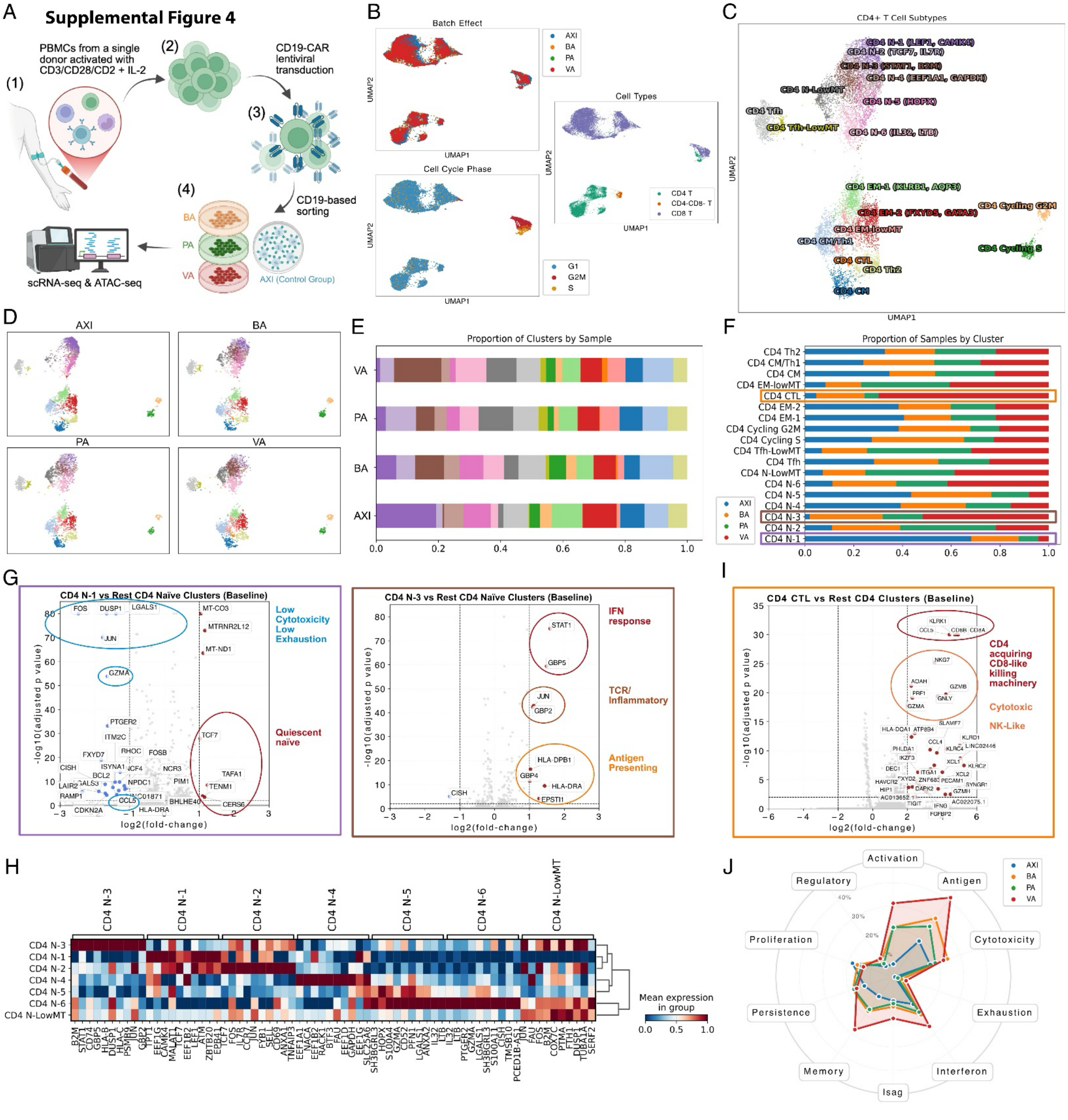


**Supplementary Figure S4.** *CD4⁺ CAR-T compartment — full cell-state panel.* Full $CD4^+$ scRNA-seq analysis of healthy-donor CD19-CAR-T cells conditioned with vehicle (AXI), BA, PA, or VA at 1 mM. **(A)** Experimental schema. **(B)** UMAPs colored by condition, cell type, and cell-cycle phase. **(C)** Annotated UMAP of 18 $CD4^+$ subclusters. **(D)** Per-condition UMAPs. **(E)** $CD4^+$ subcluster proportions by condition. **(F)** Per-cluster sample proportions with key clusters boxed. **(G)** CD4 N-1 vs N-3 baseline volcanoes. **(H)** Cluster marker heatmap. **(I)** CD4 CTL vs rest CD4 volcano. **(J)** Functional axis radar across nine T-cell programs.

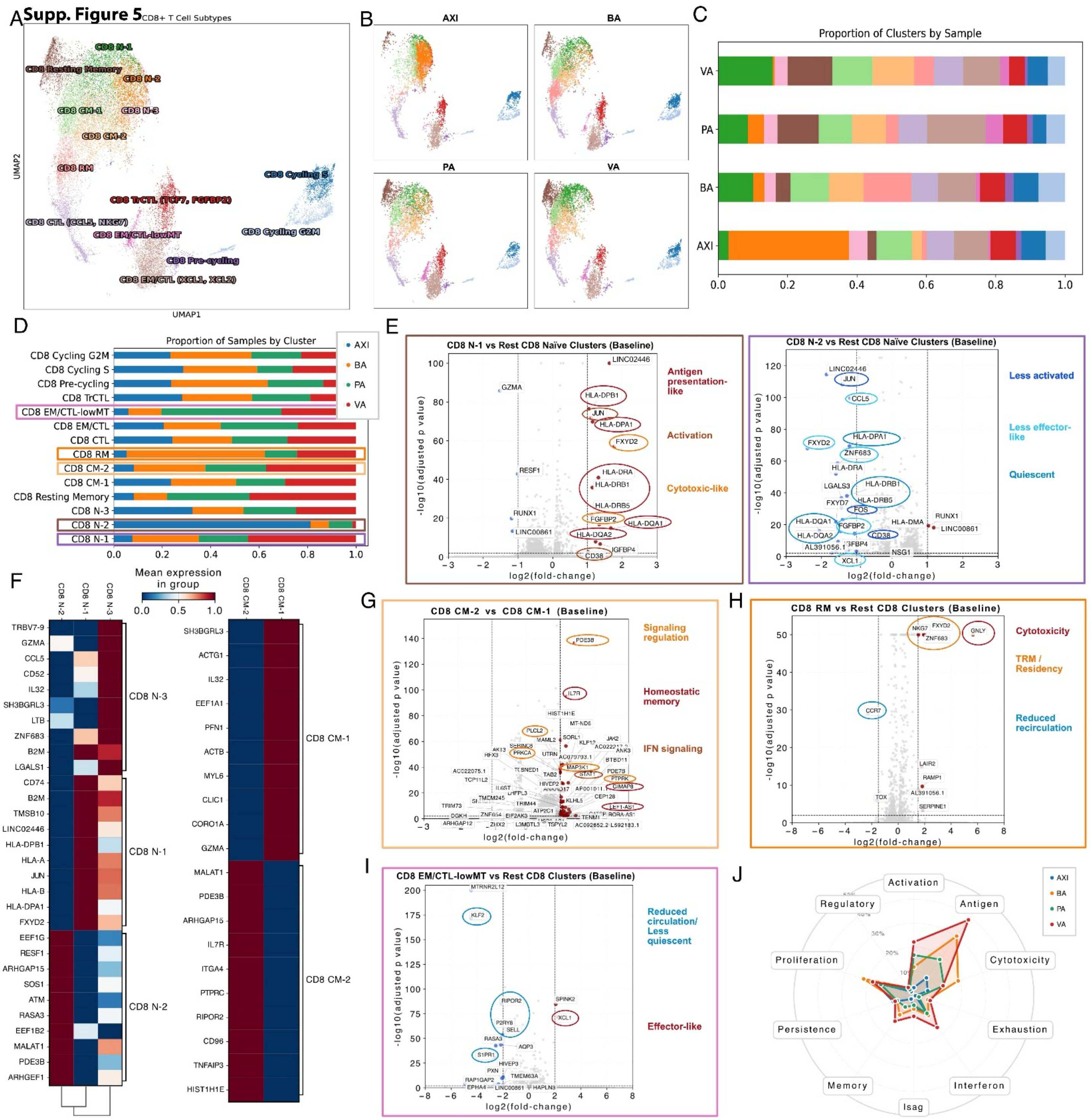

**Supplementary Figure S5.** *CD8⁺ CAR-T compartment — full cell-state panel.* CD8⁺ counterpart of S4. **(A)** Annotated UMAP of 14 CD8⁺ subclusters. **(B)** Per-condition UMAPs. **(C)** CD8⁺ subcluster proportions by condition. **(D)** Per-cluster sample proportions with key clusters boxed. **(E)** CD8 N-1 vs N-2 vs rest naïve baseline volcanoes. **(F)** Cluster marker heatmaps (CD8 N-1/N-2/N-3 and CM-1/CM-2). **(G)** CD8 CM-2 vs CM-1 volcano. **(H)** CD8 RM vs rest volcano. **(I)** CD8 EM/CTL-lowMT vs rest volcano. **(J)** Functional axis radar.

**Supplemental figure 6**

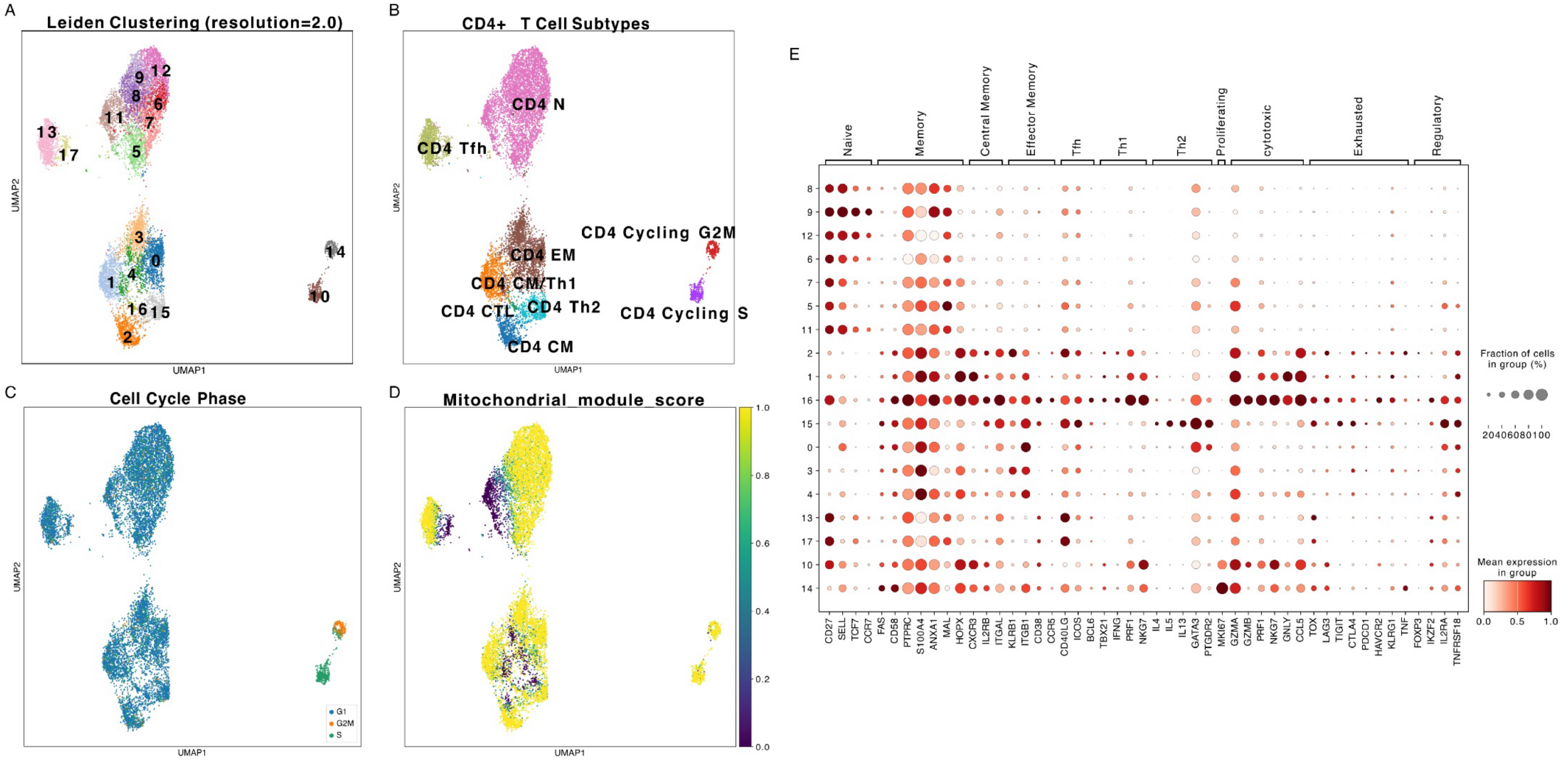


**Supplementary Figure S6.** *CD4⁺ clustering diagnostics.* **(A)** UMAP colored by Leiden clustering at resolution 2.0. **(B)** UMAP of annotated CD4⁺ subtypes. **(C)** UMAP colored by cell-cycle phase, confirming CD4 Cycling S and Cycling G2M correspond to S-phase and G2/M-phase cells. **(D)** UMAP colored by mitochondrial module score, identifying CD4 N-LowMT, CD4 Tfh-LowMT, and CD4 EM-LowMT as the low-MT subclusters. **(E)** Dot plot of canonical lineage-marker mean expression and fraction-of-cells-expressing across all 18 CD4⁺ subclusters.

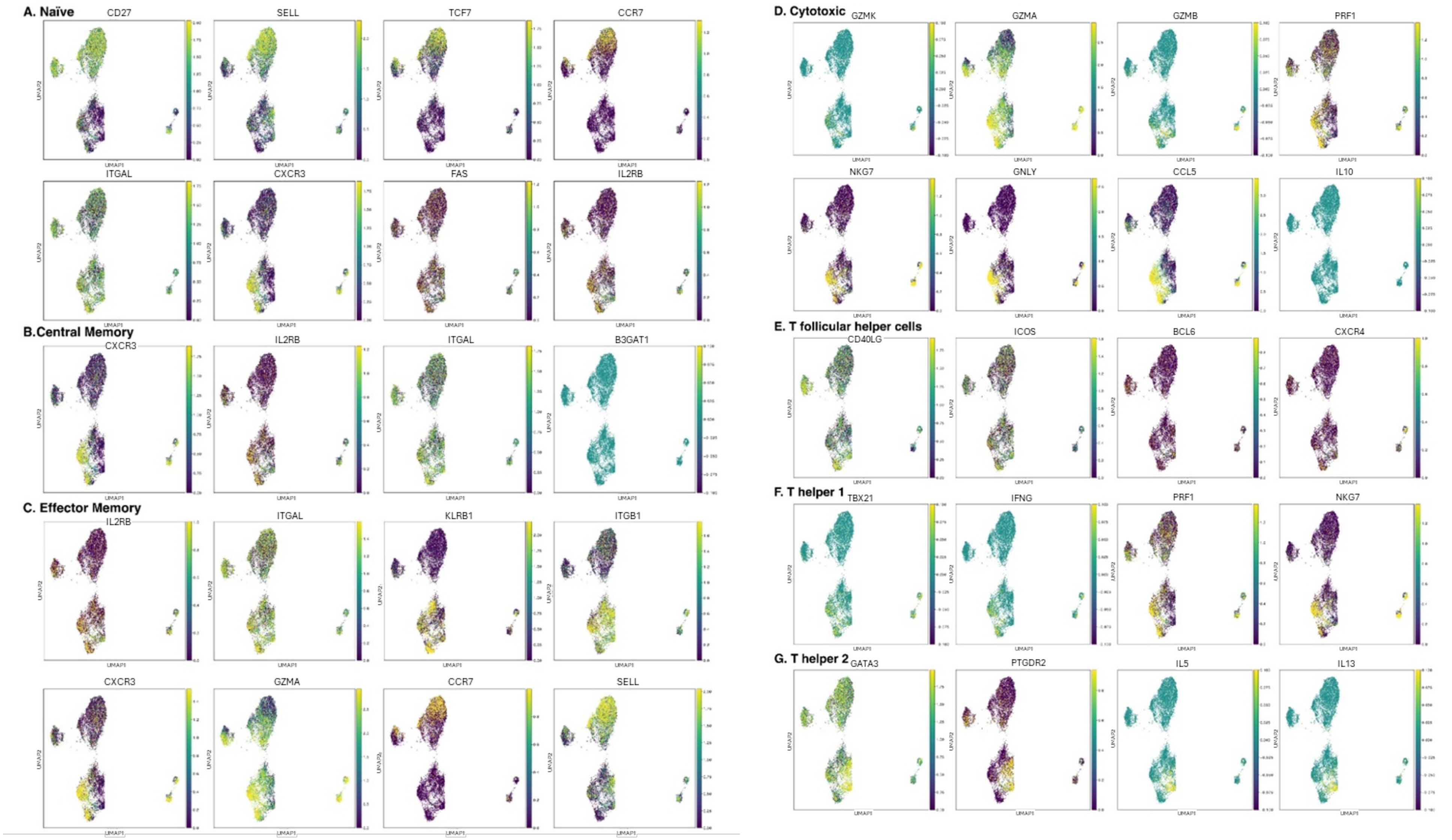


**Supplementary Figure S7.** *$CD4^+$ lineage-signature scoring per condition.* UMAP projections of canonical T-helper / cytotoxic signature scores across $CD4^+$ cells, plotted separately for each condition (AXI, BA, PA, VA; columns) and signature (rows A–G). **(A)** Naïve (SELL, CCR7, TCF7, LEF1, IL7R). **(B)** Central memory. **(C)** Effector memory. **(D)** Cytotoxic (GZMA, GZMB, PRF1, NKG7, GNLY). **(E)** T follicular helper. **(F)** Th1 (TBX21, IFNG). **(G)** Th2 (GATA3, IL4, IL13). Per-condition visualization confirms cluster-resolution lineage identity is preserved across SCFA arms.

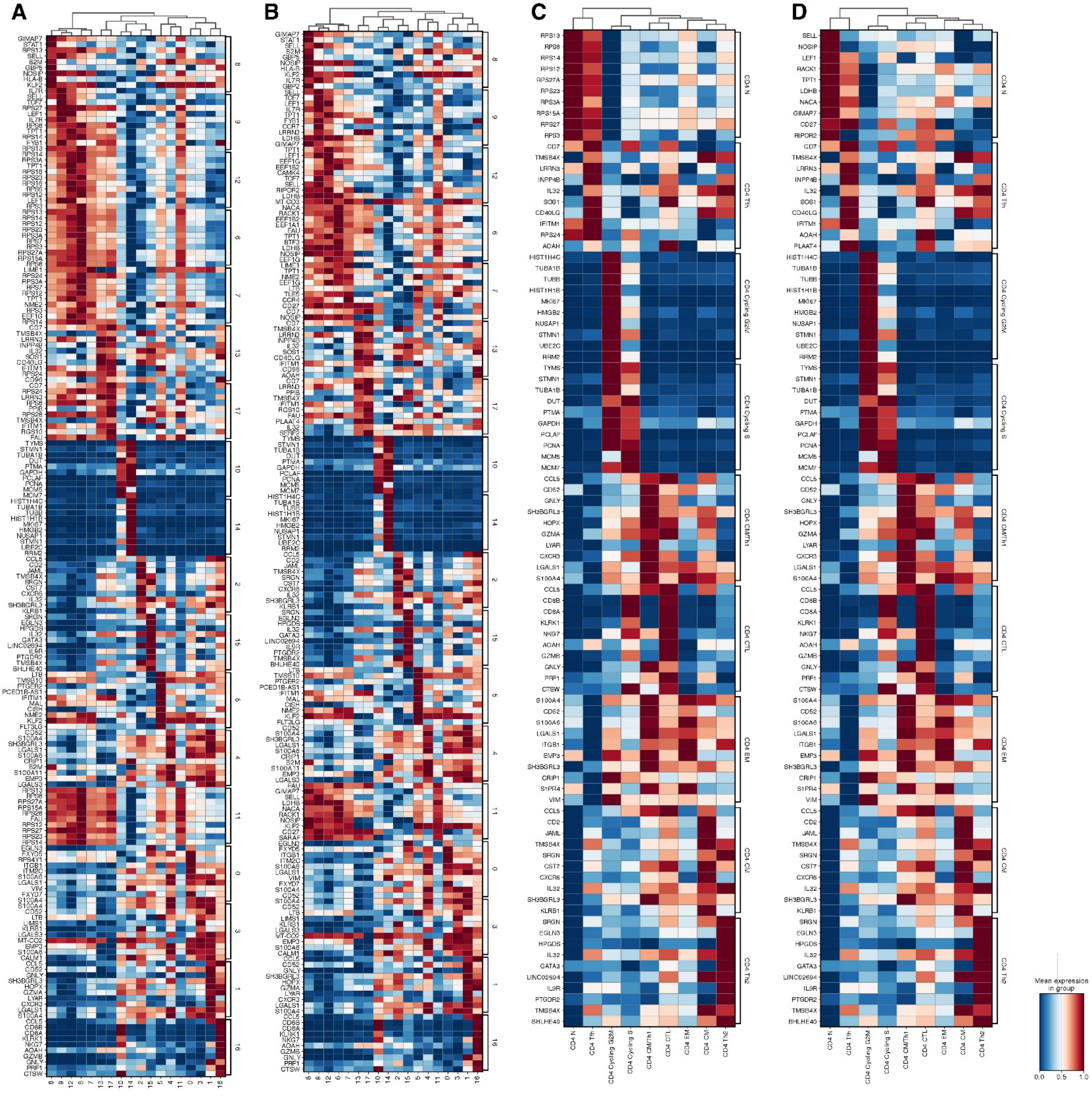


**Supplementary Figure S8.** *CD4⁺ cluster-defining DEG heatmaps.* Heatmaps of mean expression of top cluster-defining DEGs across all 18 CD4⁺ subclusters with hierarchical clustering by transcriptional similarity. **(A, B)** Complete CD4⁺ DEG reference. **(C, D)** Focused subcluster-by-subcluster DEG profiles. Color: row-normalized mean expression (blue low, red high). Naïve subclusters cluster together and separate from memory / effector / cytotoxic subclusters; CD4 N-LowMT groups with naïve states; CD4 CTL groups with CD8-lineage-marker-acquiring effector cytotoxic states.

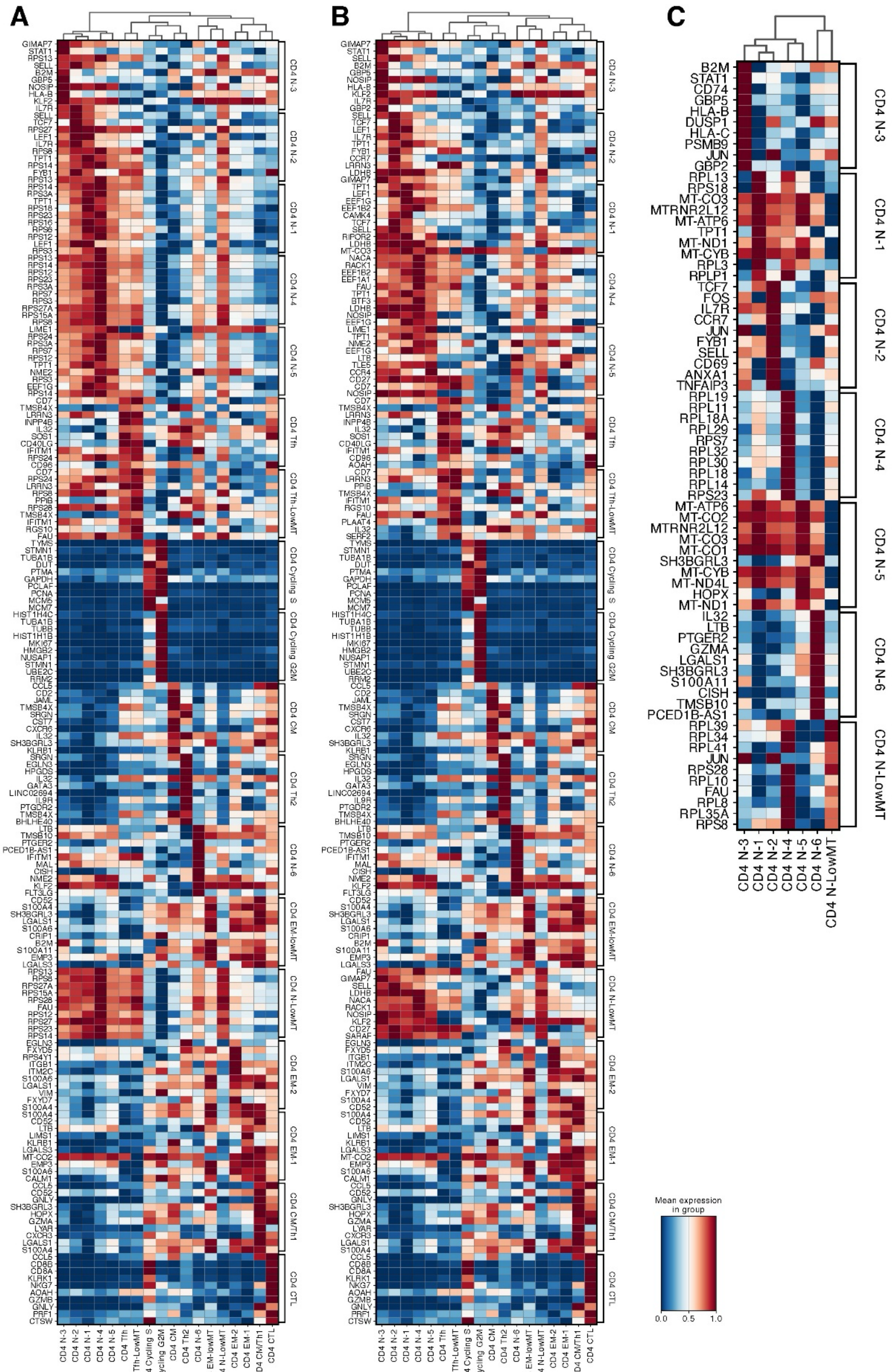


**Supplementary Figure S9.** *CD4⁺ naïve subcluster marker heatmaps.* **(A, B)** Top marker-gene expression heatmaps across CD4⁺ naïve subclusters (N-1 through N-6 plus N-LowMT) with hierarchical clustering, revealing TCF7/LEF1 high in CD4 N-1; STAT1/GBP family elevated in CD4 N-3 and N-5; mitochondrial-gene depletion in CD4 N-LowMT. **(C)** Condensed heatmap of canonical naïve-cluster-distinguishing genes (B2M, STAT1, HLA family, AP-1 family, CCR7, IL7R, ISG15, MT-encoded) defining the CD4 N-3 activated-naïve identity (AP-1 + MHC II + IFN-target).

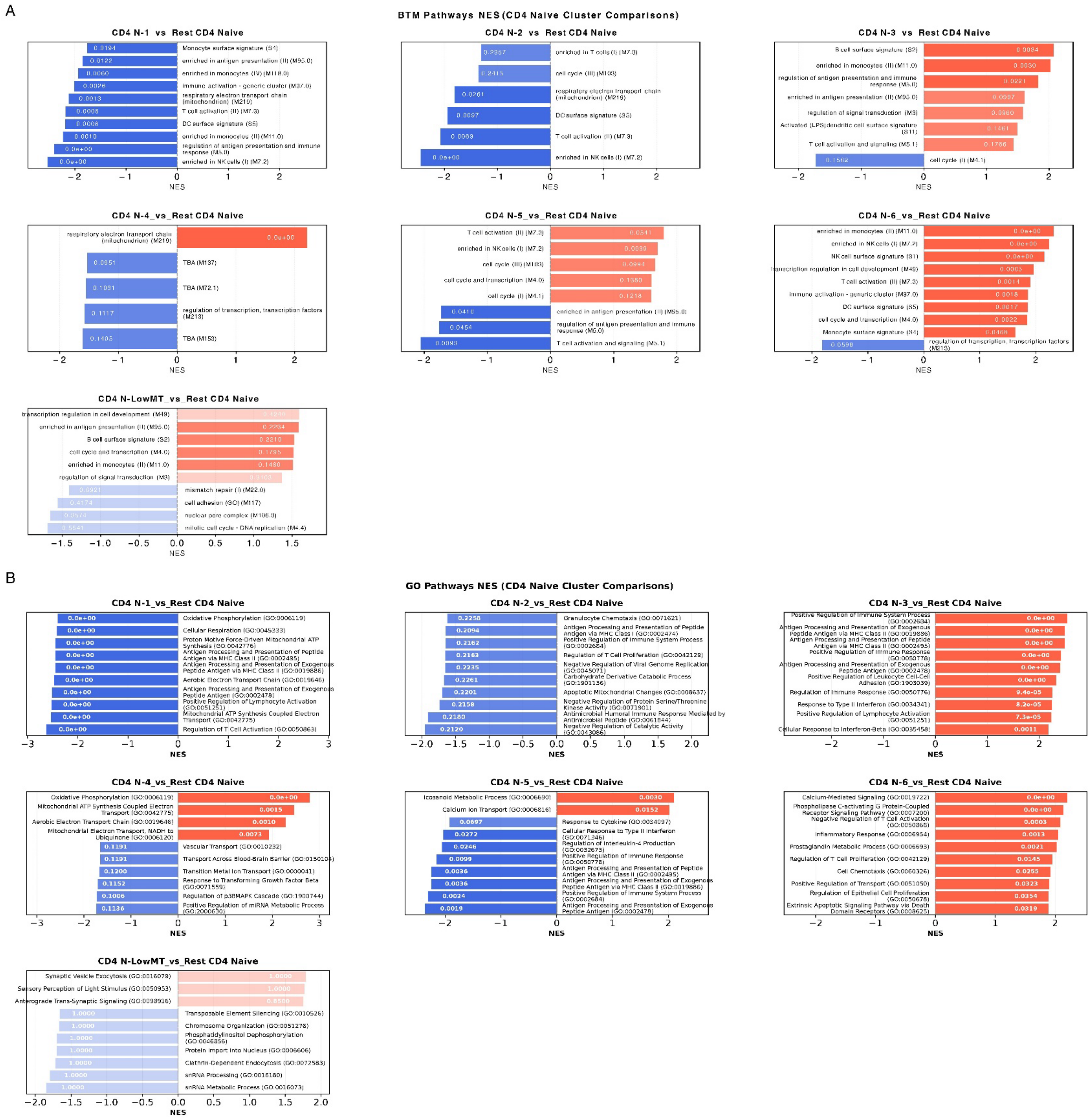


**Supplementary Figure S10.** *CD4$^+$ naïve subcluster pathway enrichment.* Per-cluster pathway enrichment for each CD4$^+$ naïve subcluster vs the remaining CD4$^+$ naïve clusters. **(A)** Blood Transcription Module (BTM) NES; CD4 N-3 and N-6 enrich for T-cell activation and MHC class II antigen processing; CD4 N-1/N-2 enrich for ribosomal/translation modules. **(B)** Gene Ontology (GO) pathway NES confirming antigen processing and presentation via MHC class II, NK/T-cell-mediated cytotoxicity, and inflammatory response as top-enriched programs in activated naïve subclusters. FDR-adjusted significance shown.

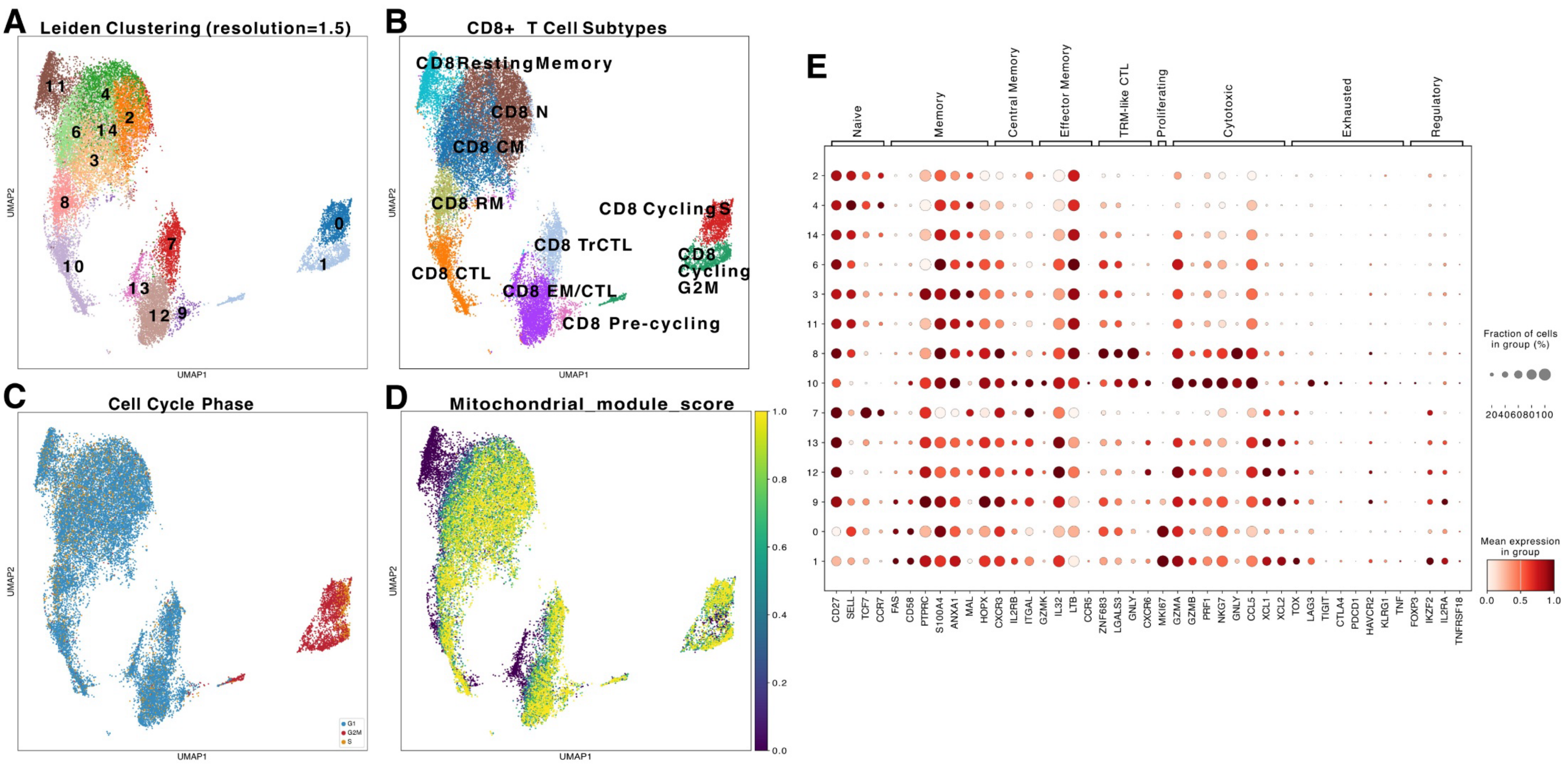


**Supplementary Figure S11.** *CD8⁺ clustering diagnostics.* **(A)** UMAP colored by Leiden clustering at resolution 1.5. **(B)** UMAP of annotated CD8⁺ subtypes (CD8 N, Resting Memory, CM, RM, CTL, TrCTL, EM/CTL, EM/CTL-lowMT, Pre-cycling, Cycling S, Cycling G2M). **(C)** UMAP colored by cell-cycle phase. **(D)** UMAP colored by mitochondrial module score, identifying CD8 EM/CTL-lowMT as the low-MT subcluster. **(E)** Dot plot of canonical CD8 lineage-marker mean expression and fraction-of-cells-expressing across all 14 CD8⁺ subclusters.

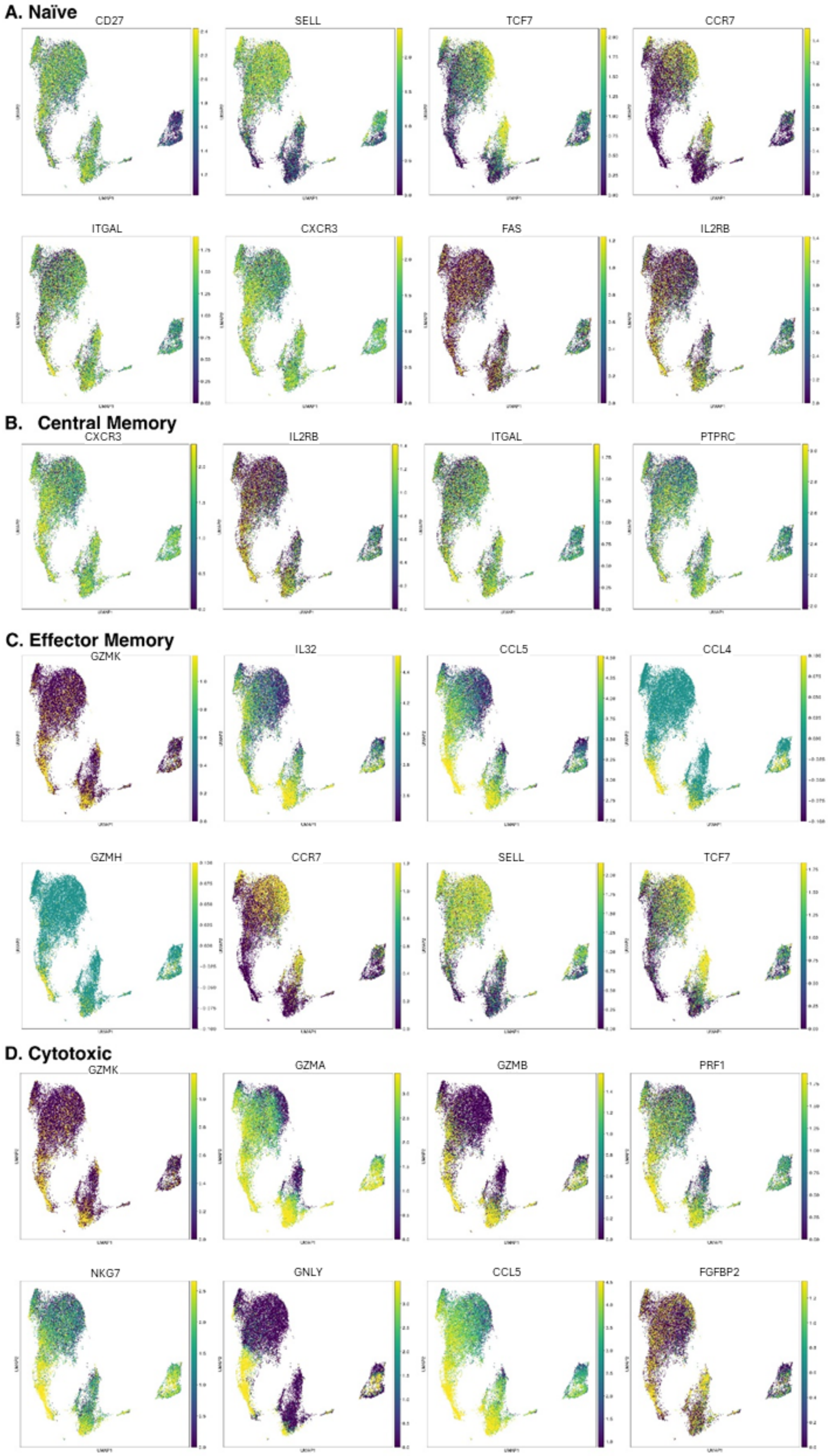


**Supplementary Figure S12.** *$CD8^+$ lineage-signature scoring per condition.* UMAP projections of canonical $CD8^+$ functional subset signatures across all $CD8^+$ cells, plotted separately for each condition (AXI, BA, PA, VA; columns) and signature (rows A–D). **(A)** Naïve, localized to CD8 N-1, N-2, N-3 and depleted under VA. **(B)** Central memory. **(C)** Effector memory, strongest enrichment under BA. **(D)** Cytotoxic, with CD8 RM preferentially enriched under VA. Per-condition visualization confirms cluster-resolution lineage identity preserved across SCFA arms.

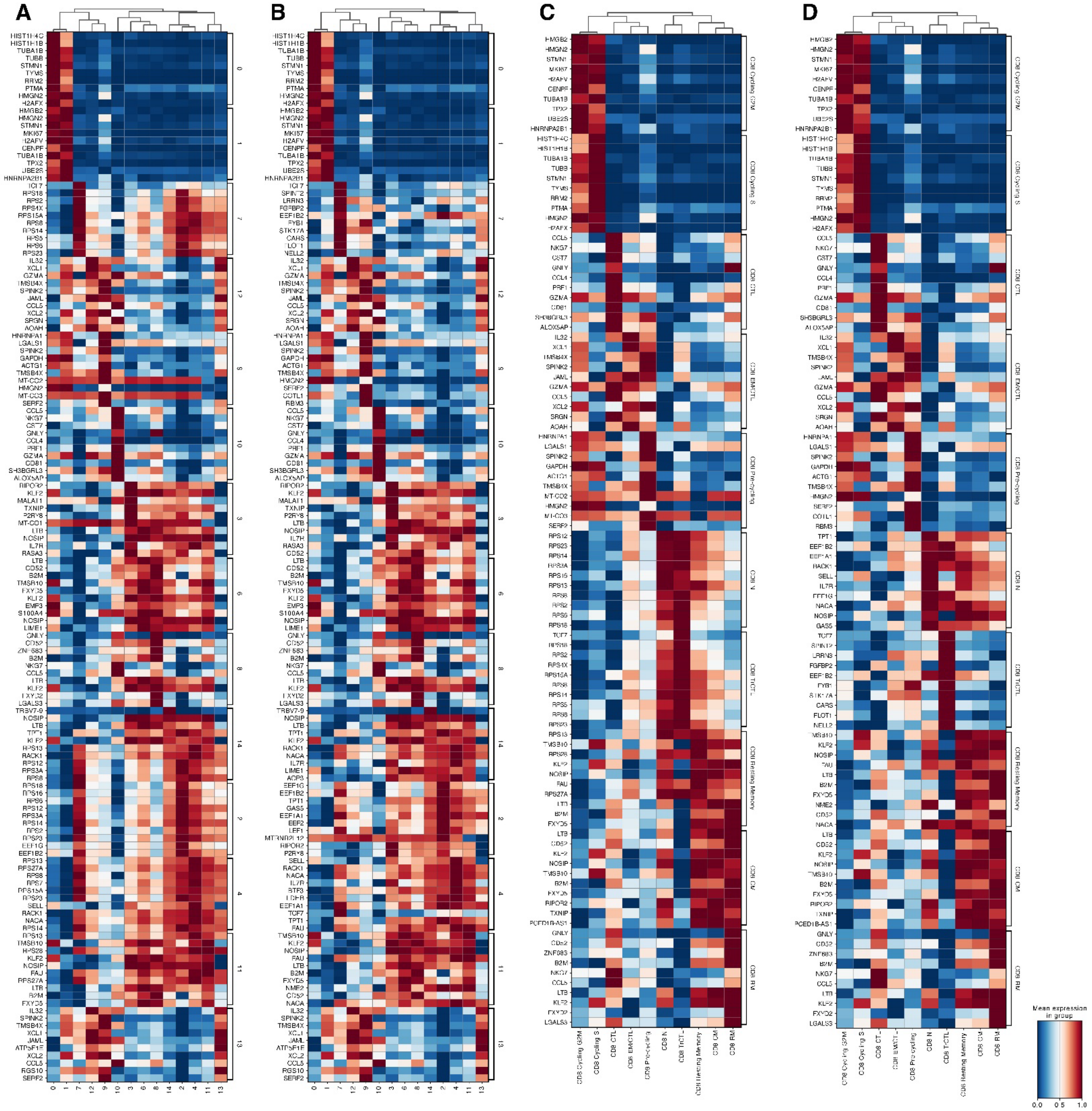

**Supplementary Figure S13.** *CD8⁺ cluster-defining DEG heatmaps.* Heatmaps of mean expression of top cluster-defining DEGs across all 14 CD8⁺ subclusters with hierarchical clustering. **(A, B)** Complete CD8⁺ DEG reference. **(C, D)** Focused subcluster-by-subcluster DEG profiles. The three naïve states (CD8 N-1, N-2, N-3) cluster together; CD8 EM/CTL and EM/CTL-lowMT group together as effector-memory cytotoxic states; CD8 RM separates from canonical effector clusters by ZNF683 (HOBIT), GNLY, NKG7 with reduced CCR7 and TOX — confirming resident-memory-like identity.

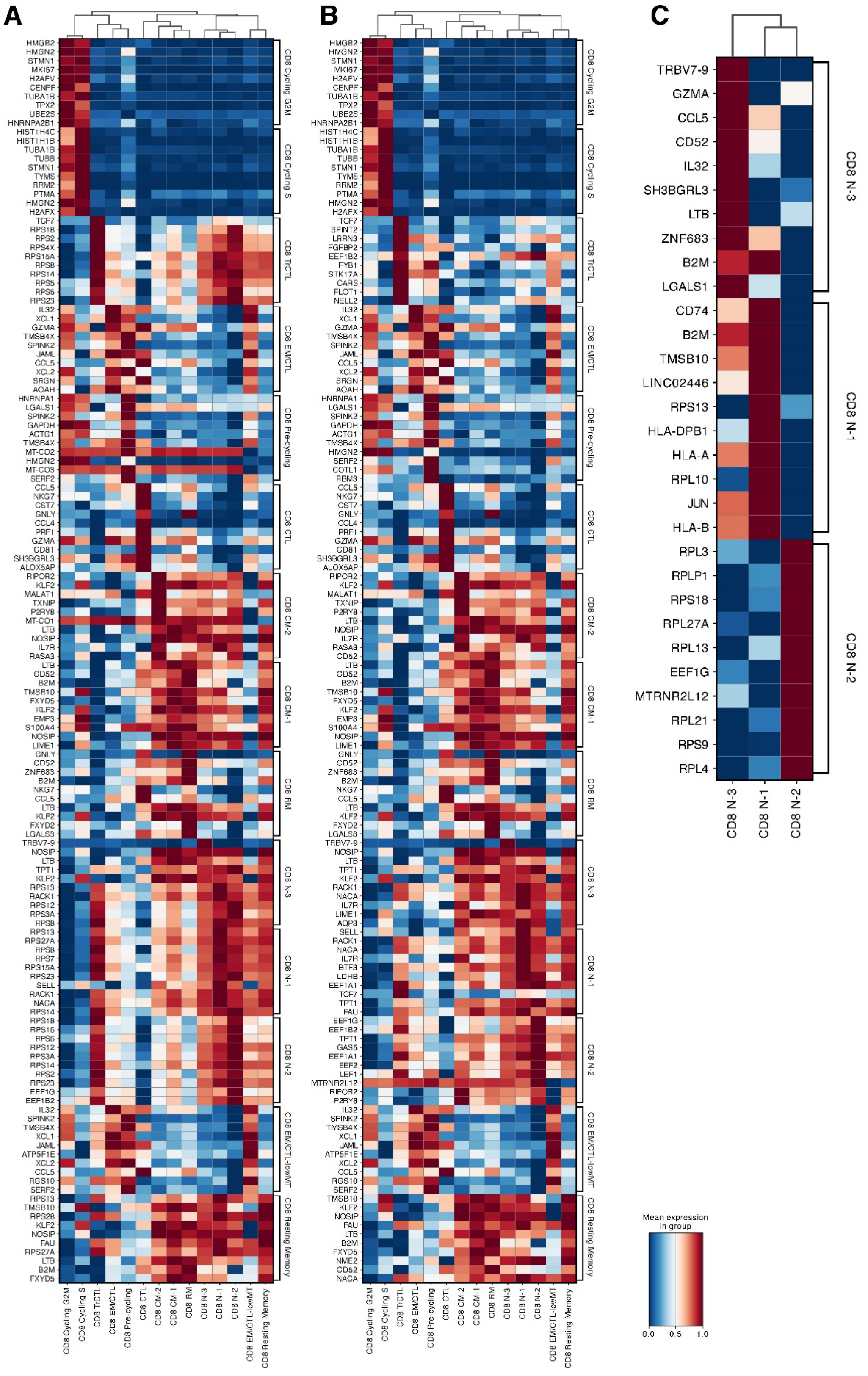


**Supplementary Figure S14.** *CD8$^+$ naïve subcluster marker heatmaps.* **(A, B)** Top marker-gene expression heatmaps across CD8$^+$ naïve subclusters (CD8 N-1, N-2, N-3) revealing CD8 N-1 as VA-enriched activated-naïve (HLA-D family, JUN, FXYD2, FGFBP2, CD38), CD8 N-2 as AXI-enriched quiescent naïve (EEF1G, RESF1, ATM, MALAT1, PDE3B), and CD8 N-3 as smaller cytotoxic-leaning naïve (TRBV7-9, GZMA, CCL5, ZNF683). **(C)** Condensed heatmap of CD8 N-1/N-2/N-3 distinguishing genes (HLA-DPB1, RPL family, B2M, JUN, ribosomal genes).

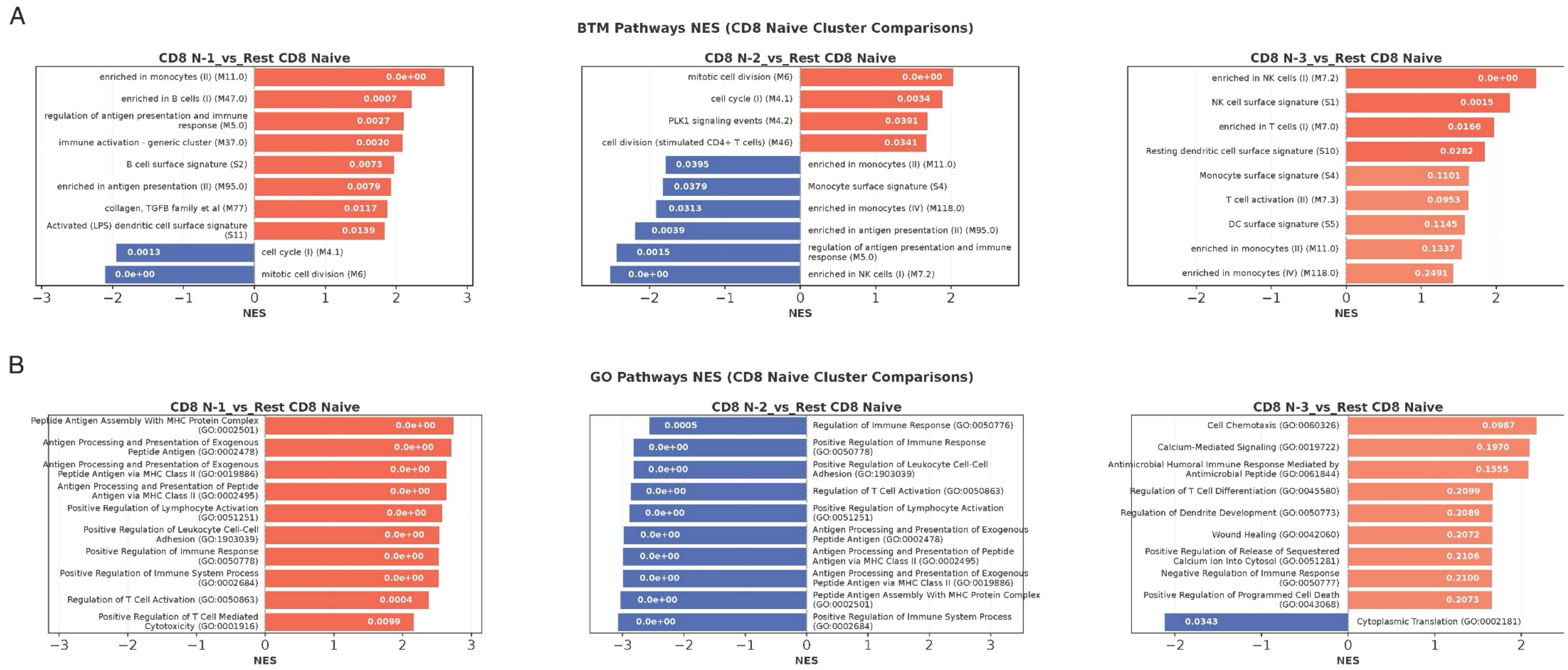


**Supplementary Figure S15.** *CD8$^+$ naïve subcluster pathway enrichment.* Per-cluster pathway enrichment for each CD8$^+$ naïve subcluster vs the remaining CD8$^+$ naïve clusters. **(A)** BTM NES; CD8 N-1 enriches for T-cell activation, MHC class II antigen processing, and regulation of antigen presentation; CD8 N-2 enriches for ribosomal/translation modules; CD8 N-3 enriches for NK-cell surface signature and cytotoxic activity. **(B)** GO pathway NES confirming activation and antigen-presentation programs in CD8 N-1, biosynthetic/ribosomal in CD8 N-2, and cytotoxic in CD8 N-3.

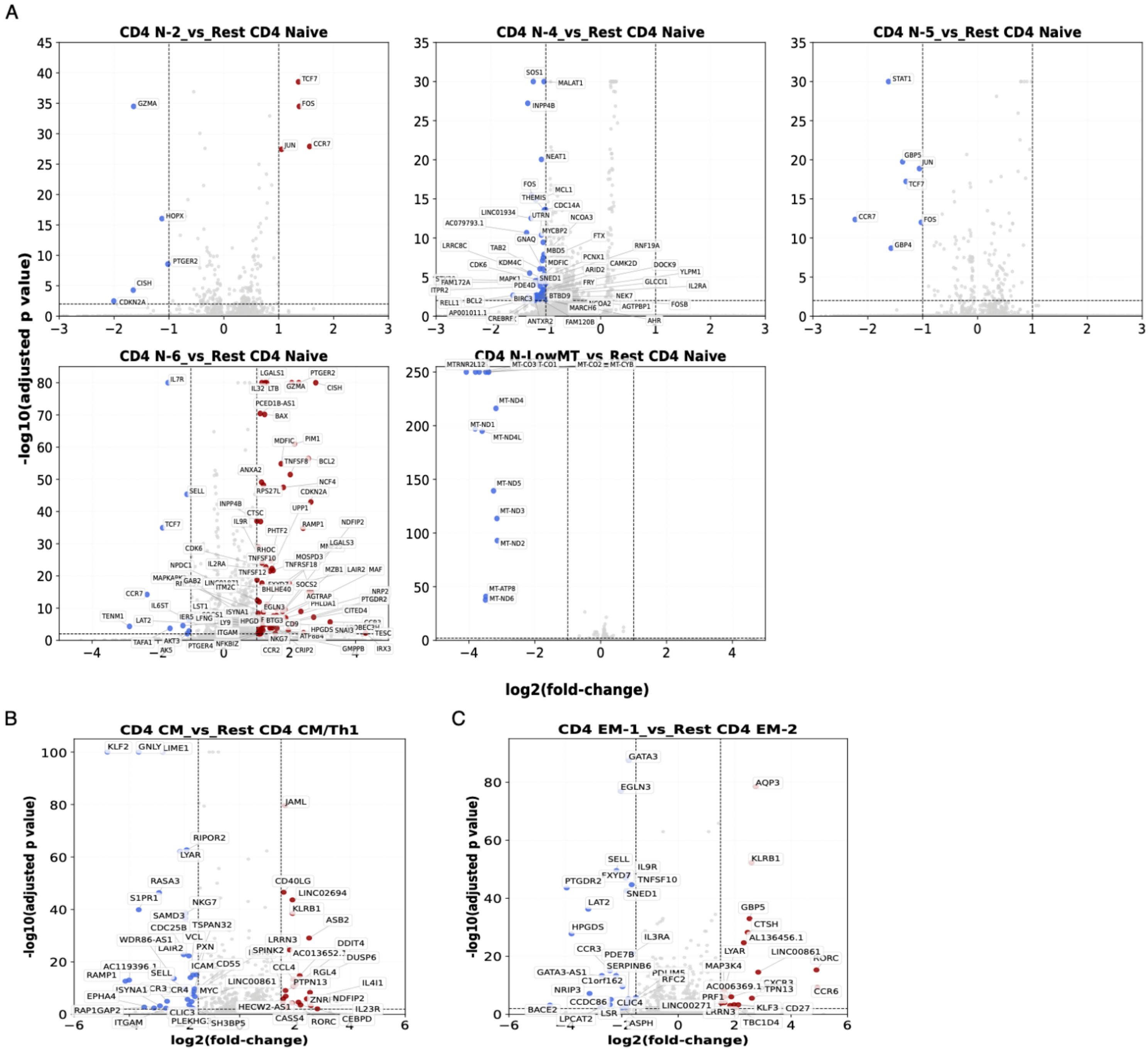


**Supplementary Figure S16.** CD4$^+$ minor-naïve and intra-family baseline volcanoes. Baseline (AXI-only) differential expression for additional CD4$^+$ subclusters not shown in main figures. **(A)** CD4 N-2, N-4, N-5, N-6, and N-LowMT each vs the rest of the CD4$^+$ naïve compartment, identifying TCF7/FOS/JUN-high quiescent (N-2), MALAT1/SOS1-defined (N-4), STAT1/GBP-defined IFN-target (N-5), activated PTGER2/LGALS1/CISH-high (N-6), and MT-genedepleted (N-LowMT) identities. **(B)** CD4 CM vs rest CD4 CM/Th1 (KLF2, GNLY high). **(C)** CD4 EM-1 vs rest CD4 EM-2 (KLRB1, RORC, AQP3 partition). Red, up; blue, down (|$\log_2$FC|>1, FDR2 & FDR2; blue, $\log_2$FC2, FDR1; blue, $\log_2$FC1; blue, $\log_2$FC<0.01.

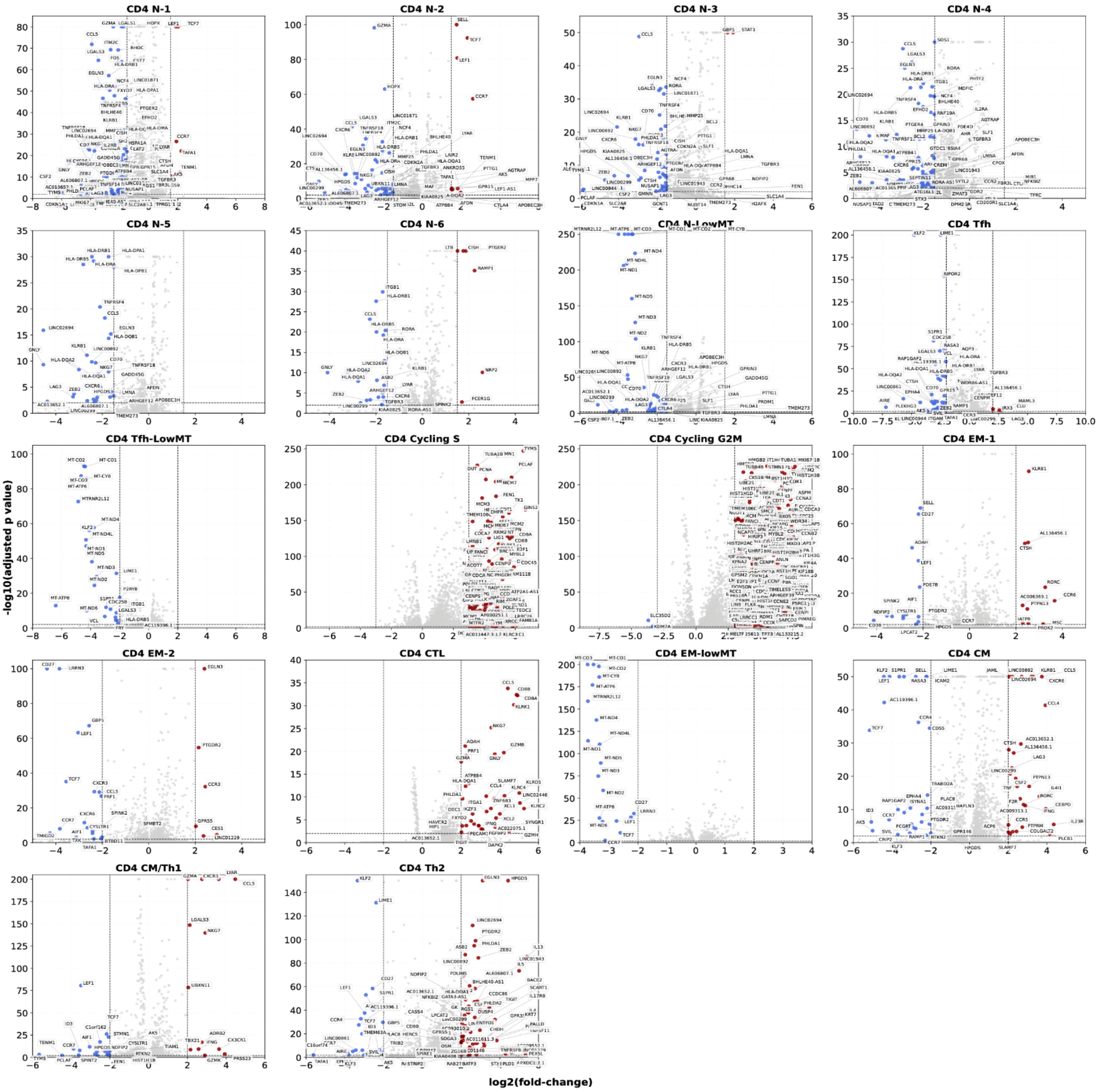


**Supplementary Figure S17. CD4$^+$ per-cluster vs rest CD4$^+$ baseline volcanoes.** Baseline (AXI-only) differential expression of each of 18 CD4$^+$ subclusters against the rest of the CD4$^+$ compartment. Panels: CD4 N-1 through N-LowMT, Tfh, Tfh-LowMT, Cycling S, Cycling G2M, EM-1, EM-2, CTL, EM-lowMT, CM, CM/Th1, and Th2. Cell-cycle clusters dominated by canonical S- and G2/M-phase genes (MKI67, HIST1H, CCNB2, TOP2A); CD4 CTL drives CCL5/CD8A/NKG7/PRF1; CD4 Th2 drives EGLN3/HPGDS/IL13/PHLDA1. Red, $\log_2$FC>2 & FDR<0.01; blue, $\log_2$FC<−2 & FDR<0.01.

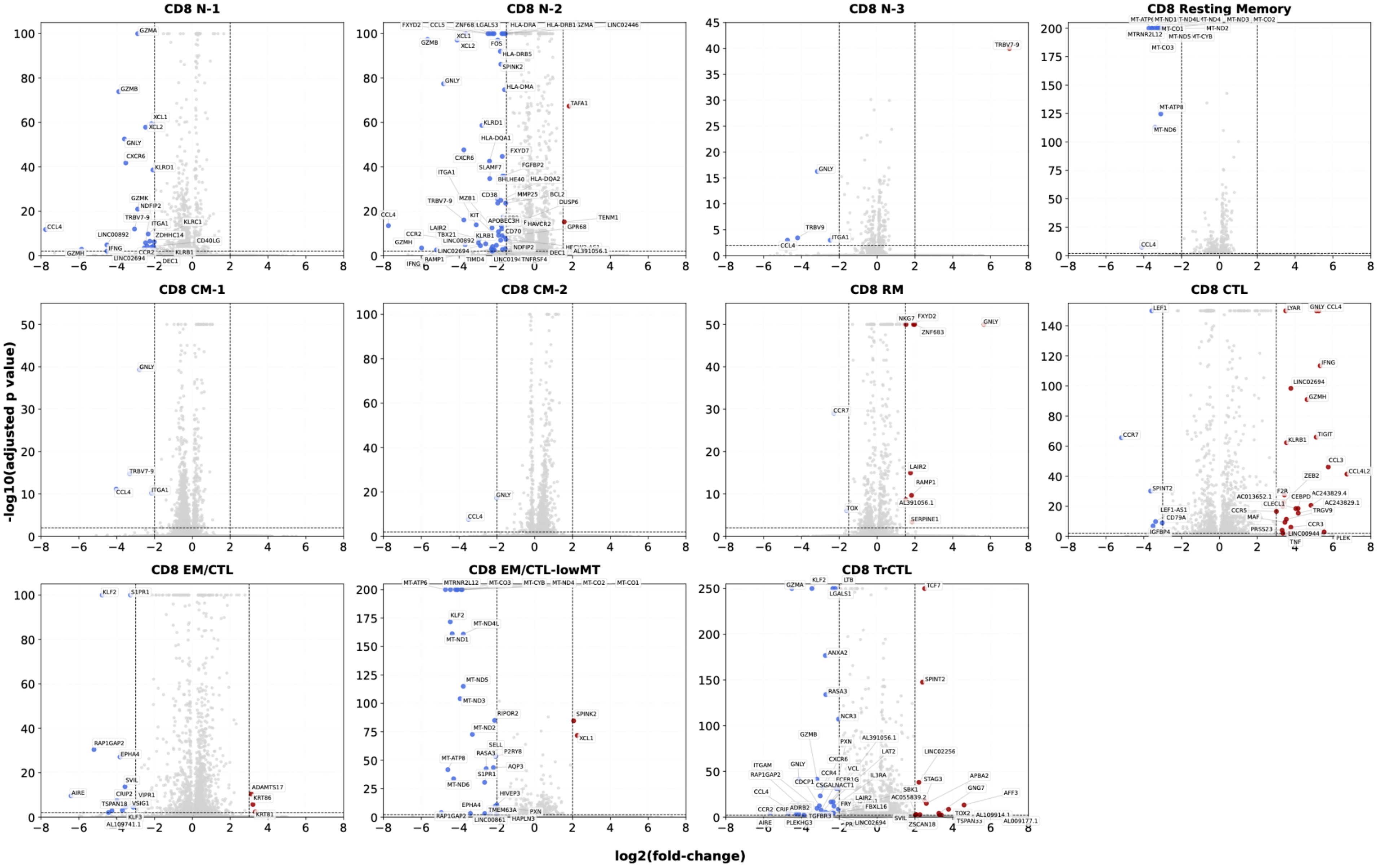


**Supplementary Figure S18. CD8⁺ per-cluster vs rest CD8⁺ baseline volcanoes.** Baseline (AXI-only) differential expression of each of 11 displayed CD8⁺ subclusters against the rest of the CD8⁺ compartment. Panels: CD8 N-1, N-2, N-3, Resting Memory, CM-1, CM-2, RM, CTL, EM/CTL, EM/CTL-lowMT, and TrCTL. CD8 N-3 driven by TRBV7-9 / cytotoxic-leaning markers; CD8 RM driven by ZNF683 (HOBIT), NKG7, FXYD2 with reduced TOX; CD8 CTL by IFNG, GZMH, KLRB1, CCL3/4/5; CD8 TrCTL separates by TCF7/SPINT2 vs GZMA/LGALS1/KLF2. Red, $\log_2FC>2$; blue, $\log_2FC<-2$ (FDR<0.01).

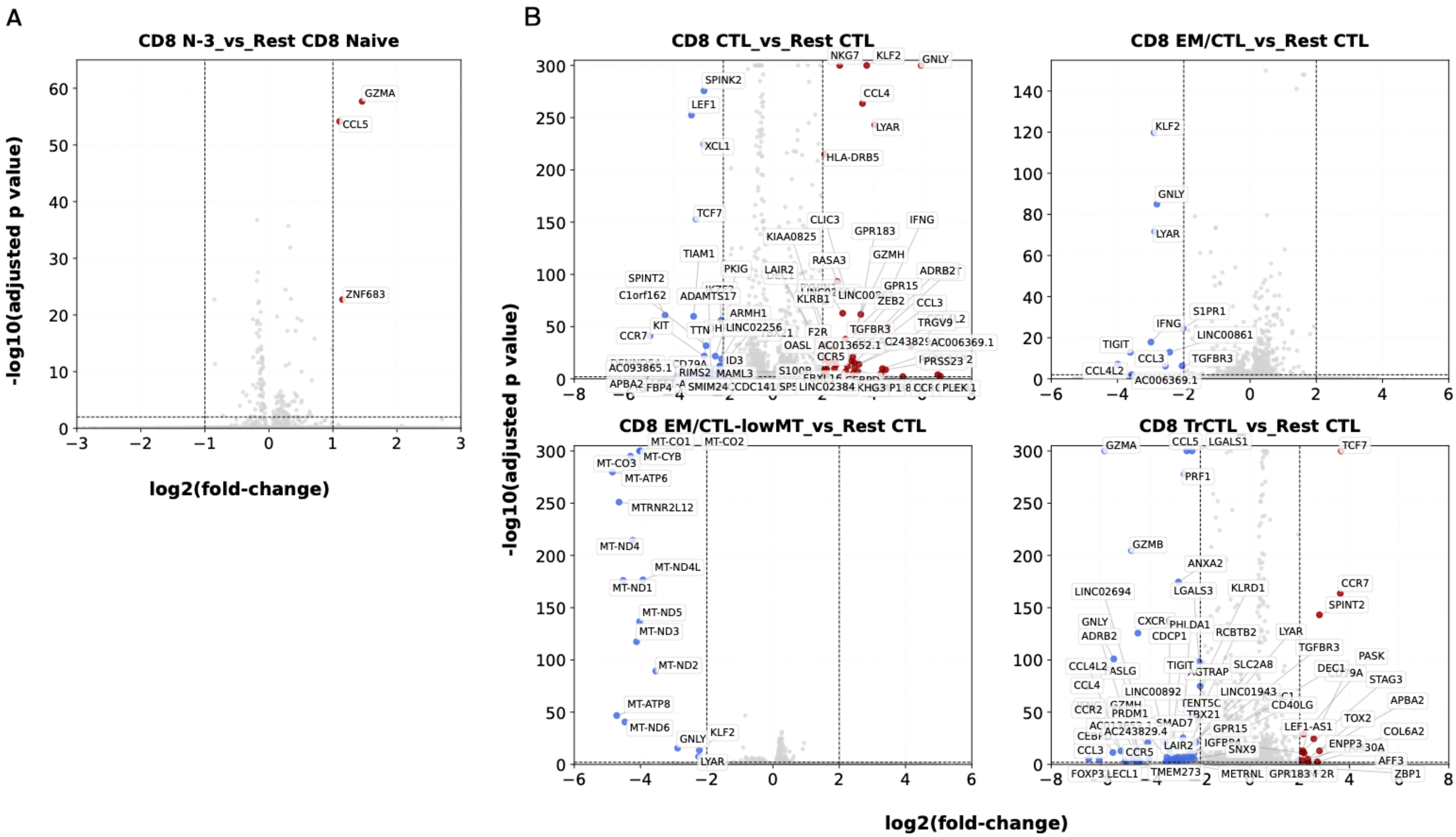


**Supplementary Figure S19. CD8⁺ N-3 and intra-CTL-family baseline volcanoes. (A)** CD8 N-3 vs rest CD8⁺ naïve subclusters, highlighting GZMA, CCL5, ZNF683 enrichment that defines this smaller cytotoxic-leaning naïve state. **(B)** Intra-CTL-family contrasts (baseline): CD8 CTL vs rest CTL (NKG7/KLF2/GNLY/CCL4/IFNG/GZMH high); CD8 EM/CTL vs rest CTL (KLF2/GNLY/LYAR down); CD8 EM/CTL-lowMT vs rest CTL (MT-gene depletion); CD8 TrCTL vs rest CTL (TCF7/SPINT2/CCR7 up, GZMA/CCL5/PRF1 down — transitional effector identity). Red, up; blue, down(|$log_2$FC|>2, FDR<0.01).

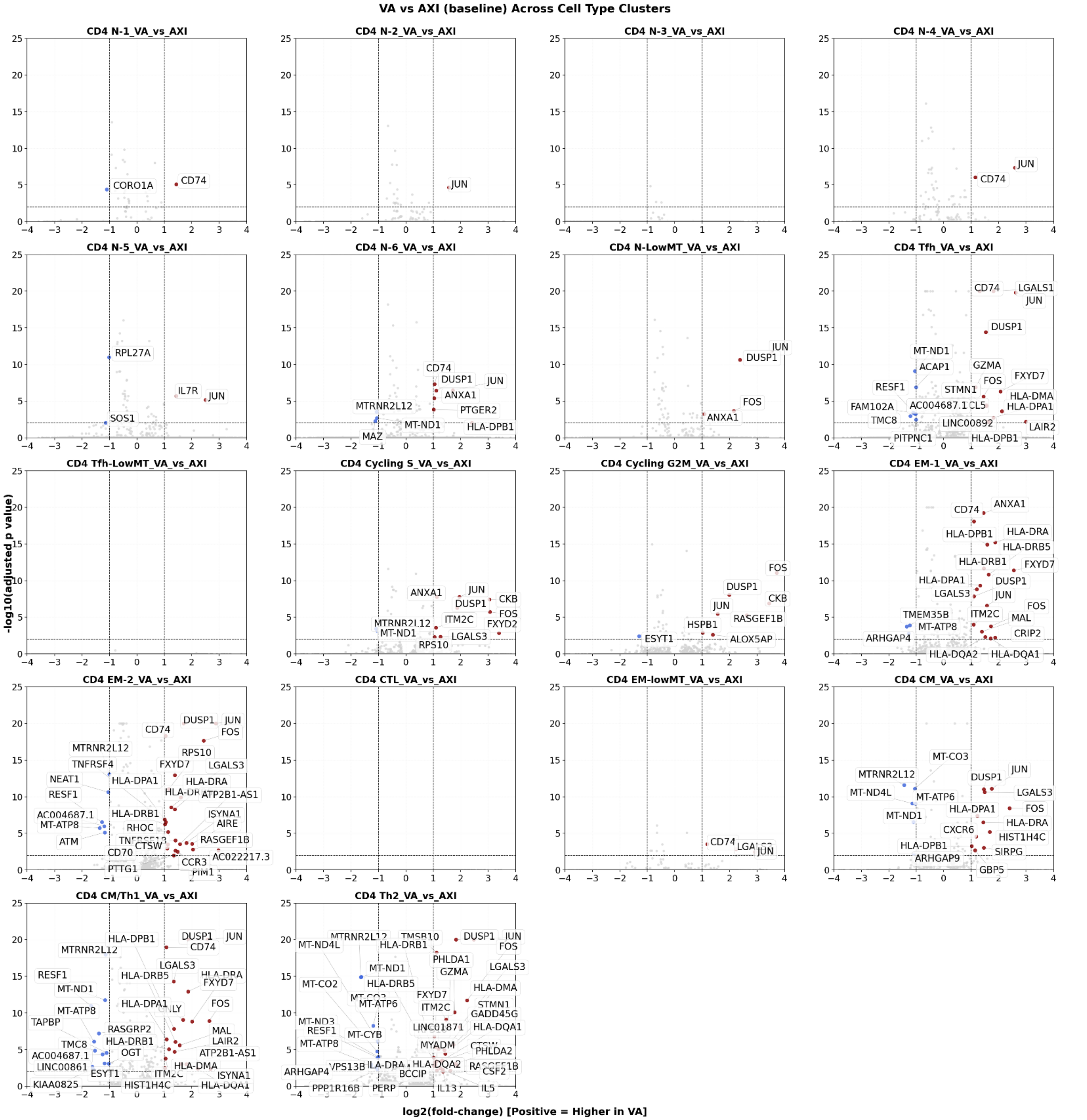


**Supplementary Figure S20. CD4⁺ VA-versus-AXI baseline per-cluster volcanoes.** Per-cluster VA-vs-AXI baseline differential expression across all 18 CD4⁺ subclusters. Positive $\log_2$FC = higher in VA. A common VA-up signature recurs across most clusters: AP-1 family (JUN, FOS, DUSP1), MHC class II (CD74, HLA-DRA, HLA-DPA1, HLA-DPB1, HLA-DRB1, HLA-DRB5, HLA-DMA), and LGALS3/ITM2C/FXYD7. Mitochondrial / RPL down-shift co-occurs in several clusters (MTRNR2L12, MT-ND1, RPL27A). Red, $\log_2$FC>1; blue, $\log_2$FC<−1; FDR<0.01. Dashed lines mark thresholds.

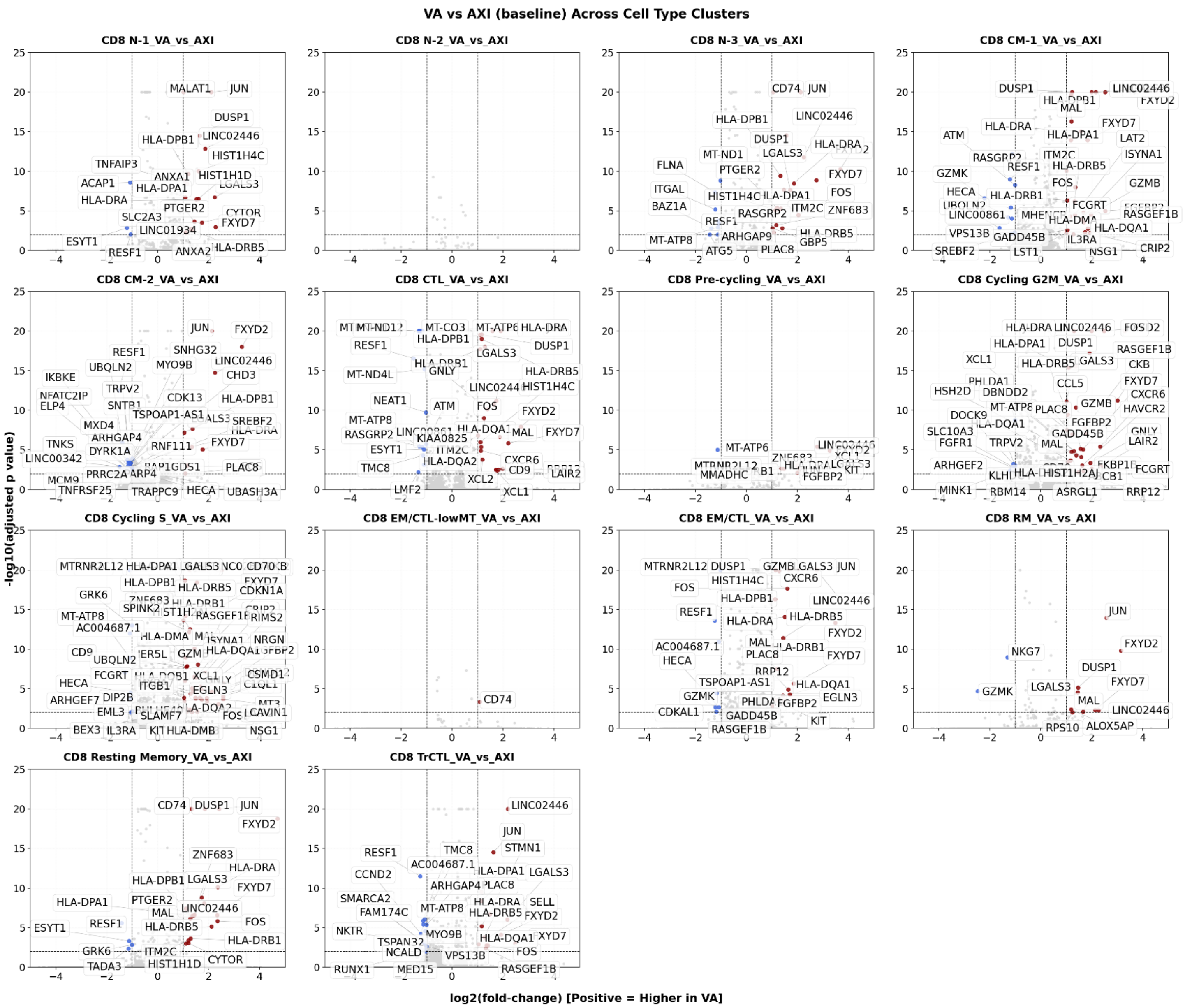


**Supplementary Figure S21. CD8$^+$ VA-versus-AXI baseline per-cluster volcanoes.** Per-cluster VA-vs-AXI baseline differential expression across all 14 CD8$^+$ subclusters. Positive $\log_2$FC = higher in VA. The same AP-1 / MHC class II VA-up program seen in CD4$^+$ recurs across CD8$^+$ clusters (JUN, FOS, DUSP1, CD74, HLA-DRA/DPA1/DPB1/DRB1/DRB5/DMA), accompanied by FXYD2/FXYD7, LINC02446, LGALS3, ZNF683 (HOBIT), and cytotoxic-program genes (GZMB, CXCR6, CCL5, HAVCR2) in CM-1, CM-2, CTL, and Cycling G2M. Red, $\log_2$FC>1; blue, $\log_2$FC<−1; FDR<0.01.
